# TEAD1 regulates ITGA1 and ITGA2 to control prostate cancer progression

**DOI:** 10.1101/2023.04.12.536554

**Authors:** Cruz Pereira Sara, Zhang Qin, Devarajan Raman, Christos Paia, Luo Binjie, Zhang Kai, Xia Jihan, Ahtikoski Anne, Vaarala Markku, Wenta Tomasz, Wei Gong-Hong, Manninen Aki

## Abstract

The extracellular matrix (ECM) undergoes significant changes during prostate cancer (PCa) progression and actively regulates PCa growth and invasion. Here, we performed a meta-analysis of PCa cohorts and found that downregulation or loss of *ITGA1* and *ITGA2* integrin genes was associated with tumor progression to metastasis and poor prognosis in PCa patients. Genomic deletion of both α1- and α2-integrins activated epithelial-to-mesenchymal transition (EMT) in benign prostate epithelial cells, thereby enhancing their invasive potential *in vitro* and converting them into tumorigenic cells *in vivo*. Mechanistically, EMT was induced by enhanced secretion and subsequent activation of autocrine TGFβ1 and nuclear targeting of YAP1. Our unbiased genome-wide co-expression analysis of large PCa cohort datasets identified the transcription factor TEAD1 as a key regulator of *ITGA1* and *ITGA2* expression in PCa cells while TEAD1 loss phenocopied the dual loss of α2- and α2-integrins in vitro and in vivo. Notably, clinical data analysis revealed that *TEAD1* downregulation or loss was associated with aggressive PCa and could synergize with *ITGA1* and *ITGA2* expression to impact PCa prognosis and progression. Altogether, our results demonstrate that loss of α1- and α2-integrins, either via deletion/inactivation of the *ITGA1*/*ITGA2* locus or via loss of *TEAD1*, contributes to PCa progression by inducing TGFβ1-driven EMT.

## INTRODUCTION

Prostate cancer (PCa) is the second most commonly diagnosed cancer and the fifth leading cause of cancer deaths in men in the western world ^1^. PCa progression is accompanied by gradual loss of epithelial organization and reorganization of the associated extracellular matrix (ECM) in the affected prostate glands. Heterodimeric integrins are the major family of ECM receptors. Integrin ligation to the ECM not only mediates adhesion but also leads to assembly of large signaling platforms that convey the microenvironmental cues to regulate cell growth, survival and migration ^2^. Several studies have implicated altered integrin functions in PCa pathogenesis ^3, 4^.

Loss of PCa cell polarity coincides with reduced expression of laminin, a central basement membrane component, and increased expression of collagen that is the major constituent of stromal ECM ^5^. Loss of β4-integrins, component of hemidesmosome-forming and laminin-binding α6β4-integrin heterodimers has been reported in PCa patients and shown to contribute to tumorigenesis ^6, 7^. α1β1- and α2β1-integrins are collagen-binding integrins that may also interact with laminins and are expressed in prostate epithelial cells ^8^. α2β1-integrins have been reported to promote tumorigenesis, particularly metastatic invasion to the bone ^9, 10^. However, other studies have observed tumor suppressive activity and/or loss of α2β1-integrin expression during PCa progression ^8, 11, 12^. To our knowledge, the potential role of α1β1-integrins or synergistic role of the neighboring *ITGA1* and *ITGA2* genes in PCa have not been addressed.

Here we have performed a meta-analysis consisting of eight PCa patient cohorts and revealed integrin signaling as the most significantly affected pathway associated with PCa progression. Remarkably, the mRNA levels of *ITGA1* and *ITGA2* genes, both encoding for collagen-binding integrin α-subunits, were markedly downregulated in PCa tumors and associated with disease progression. Therefore, we sought to study the potential role of α1β1- and α2β1-integrins in promoting tumorigenicity of PCa. Functional studies showed that both α1- and α2-integrins independently contribute to collagen adhesion and glandular morphogenesis of prostate epithelial cells. Simultaneous loss of α1- and α2-integrins in benign RWPE1 prostate epithelial cells induced cell transformation leading to increased invasive properties *in vitro* and metastatic capacity *in vivo*. This effect in α1α2-double knockout (dKO) cells was mediated by induced secretion and activation of autocrine TGFβ1 that promoted epithelial-to-mesenchymal transition (EMT) and nuclear translocation of YAP1. Transcription enhancer factor 1 (Tef-1) encoded by the TEA domain transcription factor 1 (*TEAD1*) gene, a key transcriptional co-regulator and binding partner of YAP1, was shown to bind to *ITGA1* and *ITGA2* promoter regions to induce their transcription. Loss of TEAD1 in RWPE1 cells phenocopied α1α2-dKO cells as they were transformed with increased metastatic capacity *in vitro* and *in vivo*. Taken together, our study revealed a novel mechanism where TEAD1 regulates α1- and α2-integrin expression which together exert tumor suppressive functions to maintain the epithelial phenotype by inhibiting secretion and autocrine activation of TGFβ1 in prostate epithelia. Loss of *TEAD1* and/or *ITGA1*/*ITGA2* was observed during PCa progression and was found to correlate with disease aggressiveness.

## EXPERIMENTAL PROCEDURES

### Cell culture

Benign RWPE-1 prostate epithelial cells (ATCC® CRL-11609™) were cultured in in K-SFM + Human recombinant Epidermal Growth Factor (rEGF) and Bovine Pituitary Extract (BPE) supplemented with 1% penicillin and streptomycin (all from Thermo Fisher Scientific). PCa cell line PC-3 (CRL-1435), VCap (CRL-2876), DuCap (CVCL_2025), 22Rv1 (CRL-2505), LNCaP (CRL-1740) and DU145 (HTB-81) were purchased from ATCC®,. VCaP and 293T-D10 (a kind gift from Dr. Peppi Karppinen, University of Oulu) were grown in Dulbecco’s Modified Eagle’s Medium (DMEM) (11965092, Thermo Fisher Invitrogen), for DU145 Eagle’s Minimum Essential Medium (EMEM) (30-2003, ATCC) was used. LNCaP, VCaP and 22Rv1 were grown in Roswell Park Memorial Institute Medium (RPMI 1640) (R8758, Sigma). PC-3 were grown in F12-K (30-2004, Invitrogen). DMEM, EMEM and RPMI were supplied with a final concentration of 10% fetal bovine serum (16000044, Thermo Fisher) and 1% of penicillin and streptomycin (15140122, Thermo Fisher). All cell lines were cultured in humidified incubators at 37 °C and 5% CO2.

### Antibodies and reagents

Primary antibodies are listed in Table S1. All secondary antibodies, nonspecific sheep, rabbit and mouse IgG isotype controls were purchased from Jackson ImmunoResearch. TRICT-phalloidin and DAPI were from Merck.

### Lentivirus-mediated gene knockout

Gene knockouts using the lentiviral CRISPR/Cas9-methodology were generated as described previously ^13, 14^. In short, two separate target sequences from constitutive early exons were designed for each gene (Table S2). Target sequences with no predicted off-target sites (a minimum of three mismatches with any other site in the human genome (GRCh38.p13)) were selected based on FASTA similarity search tool (EMBL-EBI). For *ITGA1*, *ITGA2* and *TEAD1*-KO1 constructs gRNA oligos with BsmBI (New England Biolabs) overhangs were ordered from Sigma-Aldrich and subcloned into lentiCRISPRv2-puro (Addgene plasmid # 52961, a gift from Feng Zhang) or lentiCRISPRv2-neo (Addgene plasmid #98292, a gift from Feng Zhang) and used for lentivirus preparation. The *TEAD1*-KO2 lentiviral construct was purchased from the Biomedicum Functional Genomics Unit at the Helsinki Institute of Life Science and Biocenter Finland at the University of Helsinki. To produce lentiviruses, 70-80% confluent 293T-D10 on CellBind® 10 cm Ø tissue culture dishes (Corning) were co-transfected with lentiCRISPR (Addgene plasmid 52961), pPAX2 and VSVg plasmids using Lipofectamine® 2000 reagent (Thermo Fisher Scientific) in Opti-MEM™ (Thermo Fisher Scientific). For infection, subconfluent cultures were incubated with virus-containing supernatants in the presence of 4µg/ml polybrene (Sigma-Aldrich , TR-1003) for 24 hours, virus was removed and culture continued for 24 hours, as described before ^13^. Cells were then trypsinized and reseeded into medium containing 3 µg/ml of puromycin to select transduced cells. Clonal cell lines were established from resistant populations. To generate double α1/α2-KO cells, clonal α1KO cells were transduced with α2-targeting lentiCRISPRv2-neo vector as described above, expanded and selected by different antibiotic selection, puromycin for α1-targeting lentiCRISPR vector and 500 µg/ml of neomycin for α2-targeting lentiCRISPR vector.

### Lentivirus-mediated gene knock-down

The lentiviral pLKO.1-based shRNA expressing vectors targeting *YAP1*, were purchased from the Biomedicum Functional Genomics Unit at the Helsinki Institute of Life Science and Biocenter Finland at the University of Helsinki. Second generation lentiviral vectors were packaged using 293T-D10 cells, as described above. Target cells were transduced sequentially two times and the transduced cell population was selected using 1 μg/ml puromycin (Sigma) as described previously ^15^.

## 3D cell culture

One hundred μls of Matrigel (Corning, #354230) was layered onto a 3.5 cm high glass bottom cell culture-dish (IBIDI, #81156) and allowed to solidify for 30 min at + 37°C (5% CO_2_). Control and α1KO, α2KO or α1α2dKO RWPE1 cells grown to confluency were trypsinized and counted. Five thousand cells per sample were resuspended into 100 μl of ice-cold Keratinocyte-SFM containing 2% of FBS and 2% (v/v) of Matrigel. Resuspended cells were seeded onto Matrigel-coated dishes and incubated for 10 min at + 37°C after which 1 ml of Keratinocyte-SFM medium containing and 2% Matrigel was added. Matrigel-containing medium was subsequently refreshed every two days. One week later, cells were fixed, stained using DAPI (Sigma, #D9542-10) and filamentous actin (Sigma, #A5441), and analyzed by using an Olympus FluoView FV1000 confocal microscope or Olympus Cellsense both equipped with 20x UPLSAPO objective (NA= 0.75).

### Immunofluorescence staining and microscopy

Cells were seeded on slides (2D culture) or on IBIDI glass bottom dishes (3D culture) and were cultured until desired confluency. Cells were fixed with 4% PFA in PBS+/+ (PBS with 0.9 mM CaCl_2_ and 0.5 mM MgCl_2_) for 15 min at room temperature for slide and 30min for IBIDI. Immunofluorescence staining was performed as previously described ^7^. Confocal images were acquired with the Zeiss LSM 780 laser scanning confocal microscope using 40x Plan-Apochromat objective (NA= 1.4) or with Olympus FluoView FV100 confocal microscope using 20x and 40x objective (NA= 0.75). Image acquisition software was ZEN (black edition, LSM 780; blue edition Cell Observer) and with Olympus FluoView viewer software, respectively.

### SDS-PAGE and western blotting

Confluent cultures were washed in PBS-/-(Gibco) and scraped into RIPA buffer: 0.5% SDS,150 mM NaCl, 10 mM Tris-HCl pH 8.0, 1% IGEPAL, 1% sodium deoxycholate with 2 mM PMSF (phenylmethylsulfonyl fluoride), 10 μg/mL aprotinin, and 10 μg/mL leupeptin. Protein concentration was calculated using BCA Protein Assay Kit (Pierce). 30 ug of protein lysate were separated on 7,5% SDS-PAGE gels and transferred onto a Protran pure 0.2 micron nitrocellulose (Perkin Elmer) for 1h30min in 20% ethanol, 0.192 M glycine and 0.025 M Tris. The membranes were blocked for 1h in 5% skimmed milk and probed with specific primary antibodies (Table S1) overnight at 4 °C. Secondary antibodies conjugated with HRP and Lumi-Light Western Blotting Substrate (Roche) were used to visualize specific protein bands. The bands were detected using Fujifilm LAS-3000 bioimaging or AZURE 600 and scientific research imaging equipment (Fuji photo film co., LTD)

### Measurement of secreted TGF-b

Ten milliliters of K-SFM with supplements was added onto confluent cultures of the different RWPE1 variants on Ø 10 cm tissue culture dishes. Forty-eight hours later conditioned medium (CM) was collected and centrifuged at 168xg for 5 min to remove cells and debris. CMs were concentrated using Amicon ® Ultra (Merk Millipore UFC503096) columns centrifuged at 101xg for 10 min. The 100x concentrated CM was analyzed for TGFβ by Western blotting.

### Adhesion assay

Control, α1KO, α2KO and α1α2dKO RWPE1 cells were seeded onto 6-well plates (Corning-Sigma) coated with 20μg/mL collagen-I (Advanced BioMatrix, 5153-A). After 30 min, plates were placed into Incucyte S3 automated imaging system (Essen Bioscience) and phase contrast image were taken for 24h with 3h intervals using Brightfield 20x objective. Posterior analysis of the attached cells was performed by counting the cells that were properly attached and the average cell count were subjected for statistical analysis.

### XTT assay

4000 cells were seeded on 96-wells plate and allowed to grow overnight. The Cell Proliferation Kit II XTT (11465015001, Roche) was used according to the manufacturer. Cell proliferation was examined at day 1, 3, 5 and 7 by XTT colorimetric assay (absorbance at 450 nm). At least three replicate wells were prepared per condition and the data were statistically analyzed with a two-tailed Student’s t test.

### Scratch wound assay

Cells were seeded into 96 well plates and allowed to grow to confluence. The scratch wound was done using the Incucyte Cell Migration Kit (Essen Bioscience 4493) and the wells were washed twice with PBS to remove detached cells. Fresh culture medium was added, and the wound areas were imaged at 3 h intervals using Incucyte Zoom (Essen Bioscience) equipped with a 20x objective. The wound area closure was analyzed using the Incucyte Zoom software (Essen Bioscience).

### Chromatin immunoprecipitation (ChIP)

The cells on tissue culture plates were cross-linked in final concentration of 1 % formaldehyde for 10 min at room temperature. Glycine was added to a final concentration of 125 mM to stop the reaction. Cell pellets were collected and snap frozen in liquid nitrogen. Pellets were then suspended in hypotonic lysis buffer (20 mM Tris-Cl, pH 8.0, with 10 mM KCl, 10 % glycerol, 2 mM DTT, and Complete protease inhibitor cocktail (Roche)) and incubated for up to 50 min to isolate the nuclei. The nuclei were washed twice with cold PBS and lysed in SDS lysis buffer (50 mM Tris-HCl, pH 8.1, with 0.5 % SDS, 10 mM EDTA, and cOmplete Protease Inhibitor). An average size of 400bp of chromatin was prepared by sonication (Q800R sonicator, Q Sonica). Seventy µl of Dynabead protein G (Invitrogen) slurry per each reaction was washed twice with blocking buffer (0.5 % BSA in IP buffer), followed by 10 h incubation with 7 µg of Tef-1 antibody against the target protein or control IgG in 1000 µl of 0.5% BSA in IP buffer (20 mM Tris-HCl, pH8.0, with 2 mM EDTA, 150 mM NaCl, 1%Triton X-100, and Protease inhibitor cocktail). Sonicated chromatin lysate (200-250 µg) was diluted into 1.3 ml of IP buffer and added onto bead/antibody complexes followed by incubation at 4 °C for at least 12 h. Next, the complex was washed once with wash buffer I (20 mM Tris-HCl, pH 8.0, with 2 mM EDTA, 0.1 %SDS, 1 % Triton X-100, and 150 mM NaCl) and once with buffer II (20 mM Tris-HCl pH, 8.0, with 2 mM EDTA, 0.1 % SDS, 1 % Triton X-100, and 500 mM NaCl), followed by two washes each with buffer III (10 mM Tris-HCl, pH 8.0, with 1 mM EDTA, 250 mM LiCl, 1 % Deoxycholate, and 1 % NP-40) and buffer IV (10 mM TrisHCl, pH 8.0, and 1 mM EDTA), respectively. Hundred ul of extraction buffer (10 mM TrisHCl, pH 8.0, 1 mM EDTA, and 1 % SDS) was added to extract the DNA-protein complexes from the beads. Proteinase K (5 µl from 20mg/ml stock) and NaCl final 0.3 M were added into the complexes incubating overnight at 65 °C to reverse the crosslinks of protein-DNA interactions. Finally, DNA was purified with MinElute PCR Purification Kit (Qiagen) and the target DNA fragments were analyzed by qPCR.

### Quantitative RT-PCR

RNA was isolated from cultured cell lines using RNeasy Mini Kit (QIAGEN), while DNA in these samples were removed by RNase-Free DNase (QIAGEN). The RevertAid reverse transcriptase Kit (Thermo Scientific) was used to synthesize cDNA from 2 ug RNA. The SYBR Select Master Mix (Applied Biosystems) was used in the Quantitative RT-PCR reactions. High specificity primers were selected for each target. Primer sequences are listed in Table S3. For the analysis of mRNA levels, each gene was analyzed at least in triplicate and the data was normalized against an endogenous GAPDH control. For ChIP-qPCR, all target primers had three technical replicates and the data were normalized to the control regions, then the relative enrichment of the target antibodies at target DNA fragment were determined by compared with the background (IgG control).

### Immunohistochemical staining

The tissue slides were deparaffinized by incubation for 1h at 55°C. To rehydrate, they were washed twice in xylene preceded 100% ethanol, 94% ethanol and 70% ethanol washes (5 min in each). Heat mediated antigen retrieval was executed by incubating the slides in boiling citric acid buffer (pH 6.0) for 10 min, then washed in PBS-/-, incubated with 3% H2O2 for 10 min and then blocked with 5% BSA in PBS-/-for 1h. After overnight incubation with primary antibodies at 4°C, the slides were washed in PBS-/- and probed for 1h with HRP conjugated secondary antibody. The positive antibody signal was revealed using DAB Substrate Kit (Abcam, ab64238). Slides were stained for 2 min in Harris hematoxylin solution followed by dehydration steps in 70%, 94% and 100% ethanol and two 5 min treatments in xylene and then imaged using Zeiss Axio Imager.M2 microscope.

### Cancer metastasis in vivo model

All animal experiments were approved by the Finnish National Animal Experiment Board (permissions ESAVI/3901/2021) and conducted in the Laboratory Animal Centre of the University of Oulu according to the principles of 3R (reduction, refinement, replacement). Intravenous tail injections were performed on 7 weeks old male IcrTac:ICR-Prkdc (SCID) mice (Taconic Biosciences A/S) as described in ^7^. In short, 1 × 10^5^ RWPE1 cell variants expressing luciferase and GFP were injected into the lateral tail vein (WT (n=6), α1KO (n=6), α2KO (n=6), α1α2dKO (n=6) and TEAD1KO (n=6)). One mouse from each group except the TEAD1KO-group died from unrelated causes during the experiment and were omitted from the analysis. Mice were monitored on a weekly basis using the IVIS Spectrum CT *in vivo* bioluminescence imaging system (PerkinElmer). Body weight was monitored every 3 days and no significant differences in body weights between the groups were observed. For analysis, the mice were injected with D-luciferin (Caliper Lifesciences) at 150mg/kg intraperitoneally (IP) and imaged 4 times at 5-minute intervals. Four weeks post injection, mice were euthanized, the blood was collected for analysis of circulating cancer cells and the lungs were dissected for analysis of luminescence and GFP signals using the IVIS system. The lungs were divided in two, one was embedded for cryosection (Fisher Healthcare Tissue-Plus O.C.T Compound, 23-730-571, Fisher & Paykel Healthcare AB, Helsinki, Finland) and one for formalin-fixed paraffin embedded (FFPE) sections. FFPE sections covering the entire lung were cut and stained with hematoxylin and eosin (HE). Slides were screened and metastatic lesions were captured using Zeiss Axio Imager M2 microscope with 40x objective. The size of micrometastatic lesions was determined based on sections were the individual lesions covered a largest area. The areas were measured using ImageJ.

### The recovery of circulating cancer cells

0,5-0,6 ml of blood from mice was immediately diluted in RBC lysis buffer (0.8% NH4Cl, 0.084% NaHCO3, 0.037% EDTA), mixed for 5 min, washed in PBS-/- and the recovered cells were seeded onto 10 cm dishes in K-SFM. After 2 days the cells were harvested by trypsin-treatment, washed in PBS-/- and GFP-positive cells were sorted using BD FACS Aria II (BD Biosciences).

### META analysis and pathway enrichment analysis

We incorporated a total eight PCa data sets from the GEO repository for conducting the meta-analysis to identify differentially expressed genes among normal prostate tissues, primary and metastatic tumors. The eight PCa data sets (GSE3933, GSE6099, GSE8511, GSE21034, GSE27616, GSE35988-GPL6480, GSE35988-GPL6848 and GSE62872 ^16–23)^ were downloaded by R package “GEOquery” with respective GEO IDs. Microarray expression data from each study was first normalized by logarithm transformation. Differential gene expression analysis was conducted on normalized expression data by R package “limma” ^24^. We applied adjustment to raw *P* values by using Benjamini & Hochberg method for multiple testing. Gene symbols were added with respective GEO Platform (GPL) identifiers. Each gene list was further filtered by removing probes corresponding to more than one gene, and unannotated probes and incomplete data was filtered out. Genes with the multiple probes was selected based on the lowest *P*-value. We used R package” MetaVolcanoR” ^25^ to incorporate multiple PCa cohorts with microarray data measured from different platforms for conducting the meta-analysis analysis. We applied both the vote counting approach that identifies differential expressed genes (DEG) for each study, and the Combining-approach that summarizes the fold change of a gene in different studies by the mean and calculates the gene differential expression *P* values using the Fisher method. We integrated results from both methods, applied a stringent cutoff by selecting differentially expressed genes with significant *P* values, and follow the same direction of being up or downregulated in at least 6 out of 8 data sets. We finally generated a comprehensive set of 1519 differentially expressed genes with 403 and 1116 up- and downregulated genes, respectively. We next used PANTHER Pathway ^26^ to investigate the underlying biological mechanisms of this DEG set in PCa and identified the integrin signaling pathway as the top altered cellular signaling cascade.

### Survival analysis

The Kaplan-Meier survival analysis was conducted to evaluate the impact of the copy number loss& deletion or low expression level of *ITGA1*, *ITGA2* or *TEAD1* on patient prognosis in multiple independent PCa data sets. Patients were stratified based on the copy number loss & deletion or the median expression level of *ITGA1*, *ITGA2* or *TEAD1*. For the investigation of the synergistic effect of *ITGA1*, *ITGA2* or *TEAD1* on patient survival, we included PCa patients with consesun triple high or low expression levels of *ITGA1*, *ITGA2* and *TEAD1*. To investigate whether *ITGA1*, *ITGA2* or *TEAD1* possess prognostic value for PCa patients with intermediate risk, we pre-stratified patients by Gleason score >=6, 7 or >=8 that represents low-, intermediate- and –high risks, respectively. Kaplan-Meier survival analysis were conducted using R package “Survival” (v. 3.2.13) ^27, 28^ and assessed by using log-rank test. Cox proportional hazards model ^29^ was applied to calculate the hazard ratio (HR) for assessing the relative risk between different patient groups.

### RNA-sequencing (RNA-seq) and differential gene expression analysis

The RNA-seq library of double knockout of ITGA1 and ITGA2, single Knockout of ITGA1 or ITGA2 generated paired-end raw sequence reads of 150 bp. Fastq files were first processed by FastQC for quality evaluation. Trimmomatic ^30^ was subsequently used on raw sequence reads for quality control. A second run of FastQC was applied on cleaned reads to guarantee high read quality. STAR (v2.7.2a) ^31^ was applied to align processed reads to the GRCh38 human genome reference by default settings. Uniquely mapped reads were then quantitated with parameters “-s no, -I gene_name” using HTSeq-count ^32^ with gene annotation information in gene transfer format (GTF) from Encode. Genes with low expression counts (<2 cumulative read count across samples) were excluded before conducting the differential expression analysis by DESeq2 (1.26.0). Genes with FDR <0.05 were identified as differentially expressed. Data normalization was processed by the Variance Stabilizing Transformation (VST) method from DESeq2. Heatmaps comparing gene expression levels across samples were generated by R package “pheatmap” (1.0.12).

### Gene Set Enrichment Analysis (GSEA)

GSEA ^33^ was used to interpret the biological functions underlying the altered gene expression profiles by the double or single knockout of *ITGA1* and/or *ITGA2* in the MSigDB database ^34^. We sorted genes in a descending order by “stat” statistics from the previously generated differential result table. The produced pre-ranked gene list was then used as input for conducting the GSEAPreranked test. Parameters were set as default except of following modifications: Enrichment statistic = “weight”, Max size (exclude larger sets) = 5000, number of permutations = 1000. The GSEA enrichment plots were produced by R packages “clusterProfiler” (3.14.3) and “enrichplot” (1.6.1) ^35^.

### The EMT pathway, EMT score and Gene expression correlation analysis

The EMT score was generated in the study carried out by Bayer et al ^36^. The authors reported that the EMT signature correlates with known EMT markers and could serve as an indicator. The EMT signature was devised with a panel of 76 representative genes, including *ANKRD22, ANTXR2, AP1M2, AXL, BSPRY, C1ORF116, KDF1, XXYLT1, CARD6, CDH1, CDH3, CDS1, CLDN4, CLDN7, CRB3, DSP, ELMO3, ENPP5, EPB41L5, EPHA1, EPN3, EPPK1, ERBB3, EVPL, F11R, FN1, FXYD3, GALNT3, GALNT5, ADGRF1, ADGRG1, GRHL1, GRHL2, HNMT, PATJ, ITGB6, KLC3, KRT19, KRTCAP3, LIX1L, TSKU, MAL2, MAPK13, MMP2, MPP7, MPZL2, TC2N, MUC1, NRP1, PPARG, PRR5, PRSS22, PRSS8, RAB25, ESRP1, RBPMS, S100A14, SCNN1A, SERINC2, SH3YL1, SHROOM3, SPINT2, SSH3, ST14, STAP2, EPCAM, TACSTD2, TGFBI, TJP3, TMC4, TMEM125, TMEM30B, TMEM45B, TNFRSF21, VIM, ZEB1*. The expression values of the EMT signature were calculated as z-score sums. The EMT pathway, consisting of 200 genes, was retrieved from the Hallmark gene sets from the MSigDB database ^33, 34^. The gene co-expression analysis was examined by the Pearson’s product-moment correlation. For the genome-wide co-expression analysis that identified *TEAD1* as one of the genes most correlating with *ITGA1* and *ITGA2*, we first investigated the expression correlation of genes in a global scale with I*TGA1* or *ITGA2* by Pearson correlation test. Gene ranks were represented by the Pearson coefficient and log10(P-value) in X- and Y-axis, respectively.

### Chromatin immunoprecipitation sequencing (ChIP-seq)

The ChIP-seq library was sequenced, which generated 150bp long single end reads. FastQC was used to check the quality of raw sequence reads. Trimmomatic ^30^ was then applied for the quality control process with following parameters: TruSeq3-SE.fa:2:30:10 SLIDINGWINDOW:5:20. Read length less than 10bp were filtered out. Bowtie2 ^37^ was then used to map the processed reads to the human genome hg38 with default settings. MACS2 ^38^ was applied for calling peaks with default settings. UCSC tools were applied to create ChIP-seq coverage signals and Integrative Genomics Viewer (IGV) ^39^ was used for visualization.

### Univariate analysis

For the univariate analysis, we investigated the association of the PCa patient overall survival, biochemical recurrence and metastasis with single, pairwise, and triple-wise combinations of gene expression levels of *ITGA1*, *ITGA2* and *TEAD1*. The z-score sum of gene expression was calculated and PCa patients were then stratified into higher and lower groups by the median expression values of the cumulative expression levels. The analysis was conducted by using R packages “tidyverse” ^40^, “tidytidbits” ^41^ and “survivalAnalysis” ^42^. Statistics were summarized and presented in forest plot.

### Meta-analysis for time-to-event outcomes

The meta-analysis for investigating the association between the ITGA1 or ITGA2 copy number loss/deletion and patient prognosis across multiple PCa cohorts was conducted using the R package “metafor” (v.3.4.0) in the R environment v.4.2.0 ^43^. The pooled HR was calculated by a fixed effect model ^44^, provided the I^2^ statistic was less than 30% or the fixed effects *P* value for the I^2^ statistic was greater than 0.10, indicating insignificant heterogeneity across studies ^45^.

### Statistical analysis

The RNA-seq, microarray or clinical data was acquired from the cBioPortal for Cancer Genomics ^46, 47^, NCBI Gene Expression Omnibus (GEO) repository ^49^, Oncomine ^48^, and literature. Microarray data for the Meta analysis was uniformly normalized by logarithm transformation (log2). All statistical analyses were carried out by R (v. 4.1.0) and RStudio (v.1.4.1106). Statistical tests applied across normal prostate tissues, primary and/or metastatic tumors were evaluated by meta-analysis, Mann-Whitney U test or Kruskal-Wallis H test depending on the context and the number of groups. The correction between gene expression data and clinical variables were assessed by Mann-Whitney U test or Kruskal-Wallis H test. The proportion of PCa patients with primary or metastatic tumors presenting the degree of number loss and deletion of *ITGA1* or *ITGA2* was examined by fisher-exact test. For investigating the synergistic effect of *ITGA1*, *ITGA2* and *TEAD1*, PCa patients were stratified to high and low groups by the median expression values of *ITGA1*, *ITGA2* and *TEAD1*, respectively. Patients with consensus high or low expressions of *ITGA1*, *ITGA2* and *TEAD1* were labeled as triple high or low group, respectively. The correlations between the synergic expression of *ITGA1*, *ITGA2* and *TEAD1* with different clinical variables including PSA, lymph node, tumor stage, and Gleason score were investigated by fisher-exact test. For results from microarray-based expression profiling, gene probes with lowest P values were selected. Samples with missing expression or patient survival data were excluded from analyses. *P* value <0.05 was considered to be statistically significant. Data are expressed as means ± SD of at least three independent experiments. Comparative data were analysed with the unpaired or paired Student’s t-test or one-way ANOVA as indicated using GraphPad Prism 8 software. The results were considered statistically significant when the p-value was less than 0.05 (*), 0.01 (**) or 0.001 (***).

## RESULTS

### α1β1- and α2β1-integrins are commonly lost or downregulated during PCa progression

The roles of androgens and AR-signaling are well-documented in metastatic PCa. However, several other cellular signaling cascades may contribute to PCa development and progression. To comprehensively dissect major signaling pathways involved in PCa, we performed a meta-analysis by incorporating eight clinical PCa expression profiling data (see **Methods**). The meta-analysis identified a total of 1284 downregulated and 610 upregulated genes over PCa tumor progression, respectively (**Fig. 1a**). Differentially expressed genes (DEGs) were subsequently subjected to PANTHER Pathway analysis for an over-representation test, which remarkably, revealed integrin signaling pathway, consisting of 191 genes, as the most significant cellular pathway deregulated during PCa progression (**Fig. 1b**). Notably, the meta-analysis identified a total 55 differentially expressed genes from the integrin signaling pathway and most of them were downregulated upon PCa development (**Fig. S1a**). Previous studies have shown that genes contributing to tumorigenesis are expected to manifest association between DNA copy number alterations and RNA expression levels ^50, 51^. We therefore sought to identify potential causal genes from the Integrin signaling pathway by the joint analysis of gene copy number alterations and expression levels. We first checked and compared the copy number profiles of differentially expressed genes from the Integrin signaling pathway in pan-cancer analysis including the TCGA PCa cohort. The result pinpointed *ITGA1* and *ITGA2* as the two genes with the highest proportions of deep loss in PCa compared to other cancer types (**Fig. 1c**).

**Figure 1.**
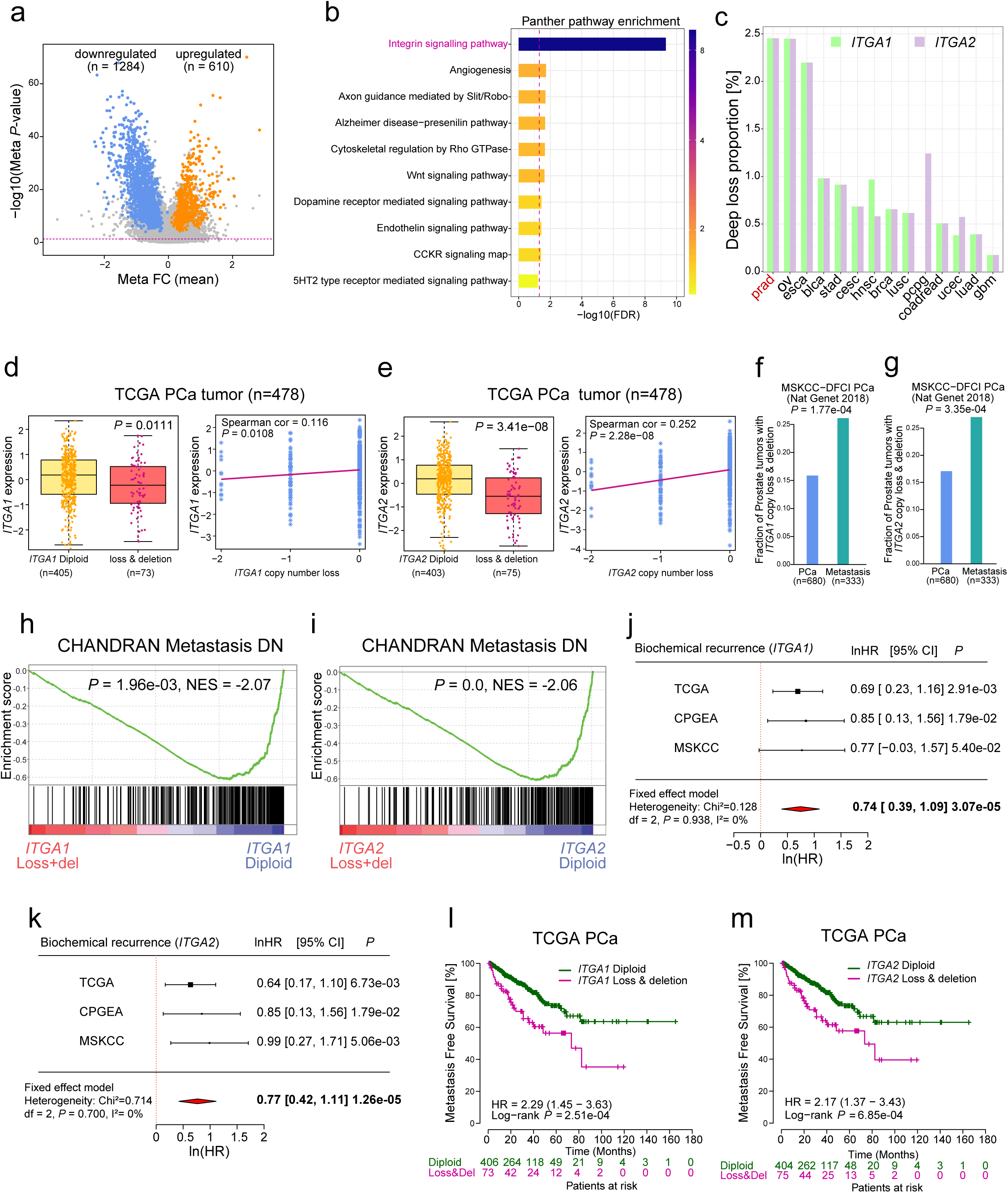
Integrin signaling pathway and the copy number loss/deletion of *ITGA1* or *ITGA2* are associated with PCa severity and prognosis. (**a**) Differentially expressed genes identified from a meta-analysis consisting of eight independent PCa cohorts. Up- and downregulated genes were highlighted in red and blue, respectively. (**b**) PANTHER pathway ontology analysis identified the Integrin signaling pathway as the most affected pathways in PCa. (**c**) *ITGA1* and *ITGA2* were identified with the highest frequencies of deep copy loss in PCa among several cancer types (ovarian - ov, esophageal - esca, bladder - blca, stomach - stca, cervical - cesc, head and neck - hnsc, breast - brca, lung - lnsc and luad, pheochromocytoma and paraganglioma - pcpg, colorectal -coadread, uterine corpus endometrial - ucec and brain - gbm). (**d-e**) *ITGA1* (**d**) or *ITGA2* (**e**) expression in prostate tumors with *ITGA1* or *ITGA2* diploid or copy number loss/deletion (left panel), and Pearson correlation between *ITGA1* or *ITGA2* mRNA expression and copy number changes (right panel). *P* values determined by the Mann-Whitney U test or from the Pearson correlation coefficient. (**f-g**) The fractions of PCa tumors harboring *ITGA1* (**f**) or *ITGA2* (**g**) copy number loss/deletion are elevated in metastasis when compared with primary PCa. *P* values were examined by the Fisher’s exact test. (**h**-**i**) Top gene set depleted in PCa tumors with *ITGA1* (**h**) or *ITGA2* (**i**) copy number loss/deletion vs diploid in the TCGA PCa cohort. NES, normalized enrichment score. (**j**-**k**) Forest plots displaying the meta-analysis of the hazard ratio estimates of *ITGA1* (**j**) or *ITGA2* (**k**) for the biochemical recurrence in multiple PCa cohorts. The horizontal error bars represent the 95% CI with the measure of center as HR. The HR and 95% CI were presented in the form of natural logarithm (ln). *P* values were calculated by the two-way Fixed-Effects Model. (**l-m**) Kaplan Meier plots indicate increased risk for metastasis in PCa patients with tumors harboring *ITGA1*(**l**) or *ITGA2* (**m**) copy number loss/deletion. *P*-values were assessed by log-rank test.

We next assessed whether the genomic alterations of *ITGA1*/*ITGA2* display any clinical impacts on PCa, and thus investigated potential correlation between *ITGA1*/*ITGA2* copy number loss and their expression levels in human prostate tumors. The results showed that *ITGA1*/*ITGA2* expression levels were dramatically decreased in PCa patient tumors with *ITGA1*/*ITGA2* copy number loss/del in two independent clinical PCa cohorts (**Fig. 1d-e** and **Fig. S1b-c**). Next, we examined whether *ITGA1*/*ITGA2* copy loss directly correlates with PCa metastasis in two independent cohorts of PCa patients with primary and metastatic tumors. This analysis revealed that the proportion of *ITGA1*/*ITGA2* copy number loss/del was substantially higher in PCa patients with metastasis compared to patients with localized tumors (**Fig. 1f-g** and **Fig. S1d-e**). The finding suggests a potential functional relevance of *ITGA1*/*ITGA2* copy loss/del in PCa carcinogenesis and tumor progression. We hence conducted a gene set enrichment analysis (GSEA) using the genome-wide expression profiling data in the TCGA cohort to identify gene sets that are enriched in phenotypes of diploid and *ITGA1*/*ITGA2* loss/del, respectively. The results demonstrated that the “CHANDRAN Metastasis DN”, a gene signature representing genes downregulated in metastatic vs nonmetastatic prostate carcinoma, was identified as the top enriched gene set across all gene sets depleted in PCa tumors with *ITGA1*/*ITGA2* loss/del compared to diploid tumors (**Fig. 1h-i**). Interestingly, we also found that several cell cycle-related pathways including E2F targets and G2M checkpoint were highly enriched in PCa tumors with *ITGA1*/*ITGA2* loss/del **(Fig. S1f-i**), further implicating the functional role of *ITGA1*/*ITGA2* loss/del in PCa tumorigenesis and metastasis.

We further evaluated the clinical impact of copy number alterations of *ITGA1*/*ITGA2* on PCa progression and therefore conducted the Kaplan-Meier survival analysis to examine the association of *ITGA1*/*ITGA2* copy number loss/del and patient prognosis in multiple independent PCa cohorts. The results displayed that PCa patients with *ITGA1*/*ITGA2* copy number loss/del were associated with shorter biochemical recurrence-free survival in three distinct PCa cohorts (**Fig. S1j-o**). To validate the robustness of the association, we additionally carried out a meta-analysis to systematically review, interpret and estimate the overall association between *ITGA1*/*ITGA2* copy number loss/del and PCa patient survival outcomes. The result demonstrated that PCa patients with *ITGA1* copy number loss/del was significantly associated with shorter biochemical recurrence-free survival (*P*=3.07e-05, lnHR= 0.74; 95% CI: 0.39–1.09). Similar association was observed for PCa tumor with *ITGA2* copy number loss/del (*P*=1.26e-05, lnHR= 0.77; 95% CI: 0.42– 1.11) (**Fig. 1j-k**). Having demonstrated the high proportion of *ITGA1*/*ITGA2* copy loss/del in metastatic PCa tumors overrepresenting metastasis pathways (**Fig. 1f-i**), we proceeded to investigate the effect of *ITGA1*/*ITGA2* copy number loss/del on patient metastatic progression outcomes. The result strongly indicated that PCa patients with *ITGA1*/*ITGA2* copy number loss/del were associated with increased metastatic risks (**Fig. 1l-m**). Taken together, these data demonstrated that genomic loss/depletion of *ITGA1*/*ITGA2* is accompanied with their decreased expression and is significantly associated with metastatic PCa progression, thereby suggesting a potential prognostic value for the *ITGA1/ITGA2*-status in PCa risk prediction.

Given that the expression levels of *ITGA1*/*ITGA2* were downregulated in PCa tumors with copy number loss/del (**Fig. 1d-e** and **Fig. S1b-c**), and that *ITGA1*/*ITGA2* genomic alterations in turn demonstrated clinical impacts on PCa development and progression (**Fig. 1j-m** and **Fig. S1j-o**), we next explored whether lower mRNA levels of *ITGA1*/*ITGA2* correlated with human PCa progression in the clinical setting and thus analyzed multiple independent PCa cohorts. The results indicated that *ITGA1*/*ITGA2* were reproducibly found to be downregulated in prostate tumor tissues compared to normal prostate glands (**Fig. 2a-d**, **Fig. S2a-c** and **Fig. S2e-g**). In particular, we observed that the mRNA levels of *ITGA1*/*ITGA2* were significantly decreased in metastatic PCa samples (**Fig. 2c-d**, **Fig. S2b-d** and **Fig. S2f-h**). To further validate this finding, we analyzed a tissue microarray consisting of matched normal vs. cancer samples our inhouse cohort of 242 PCa patients (**Table S6**). Four tissue cylinders were picked from each patient sample (two from normal/benign areas and two from carcinoma regions). In normal/benign tissue, the most intense α1-integrin expression was seen in the stromal cells of the prostate and low levels were detected in the basal epithelium whereas α2-integrin was highly expressed by basal epithelial cells and moderately by luminal epithelial cells (**Fig. 2e**). The analysis of tumor tissues confirmed a significant downregulation of α2-integrins in tumor lesions (**Fig. S3a, c**). Curiously, α1-integrin expression was significantly reduced in the tumor stroma (**Fig. S3a-b**). These results suggest a potential role of *ITGA1*/*ITGA2* in advanced prostate tumors. Thus, we next investigated the association of *ITGA1*/*ITGA2* expression with PCa patient prognosis by conducting the Kaplan-Meier analysis. The results showed that the patient group with tumors expressing lower levels of *ITGA1*/*ITGA2* had increased risk of biochemical relapse and metastasis in multiple independent PCa cohorts (**Fig. 2f-i**, **Fig. S2i-j** and **Fig. S2l-m**). Notably, we also observed that PCa patients with lower *ITGA1*/*ITGA2* mRNA levels experienced shorter time trends for overall survival (**Fig. S2k, n**). Additionally, we analyzed PCa expression data with clinical features, finding that *ITGA1*/*ITGA2* mRNA levels were significantly lower in tumors with high tumor stage, Gleason score, lymph node and elevated PSA levels in multiple distinct patient cohorts (**Fig. 2j-q** and **Fig. S2o-s**). Taken together, these results demonstrate prognostic values of *ITGA1*/*ITGA2* loss/deletion and downregulation upon PCa progression to metastatic stage, suggesting that *ITGA1*/*ITGA2* are new biomarkers that could distinguish aggressive disease and might possess potential tumor suppressive roles of *ITGA1*/*ITGA2* in PCa development.

**Figure 2.**
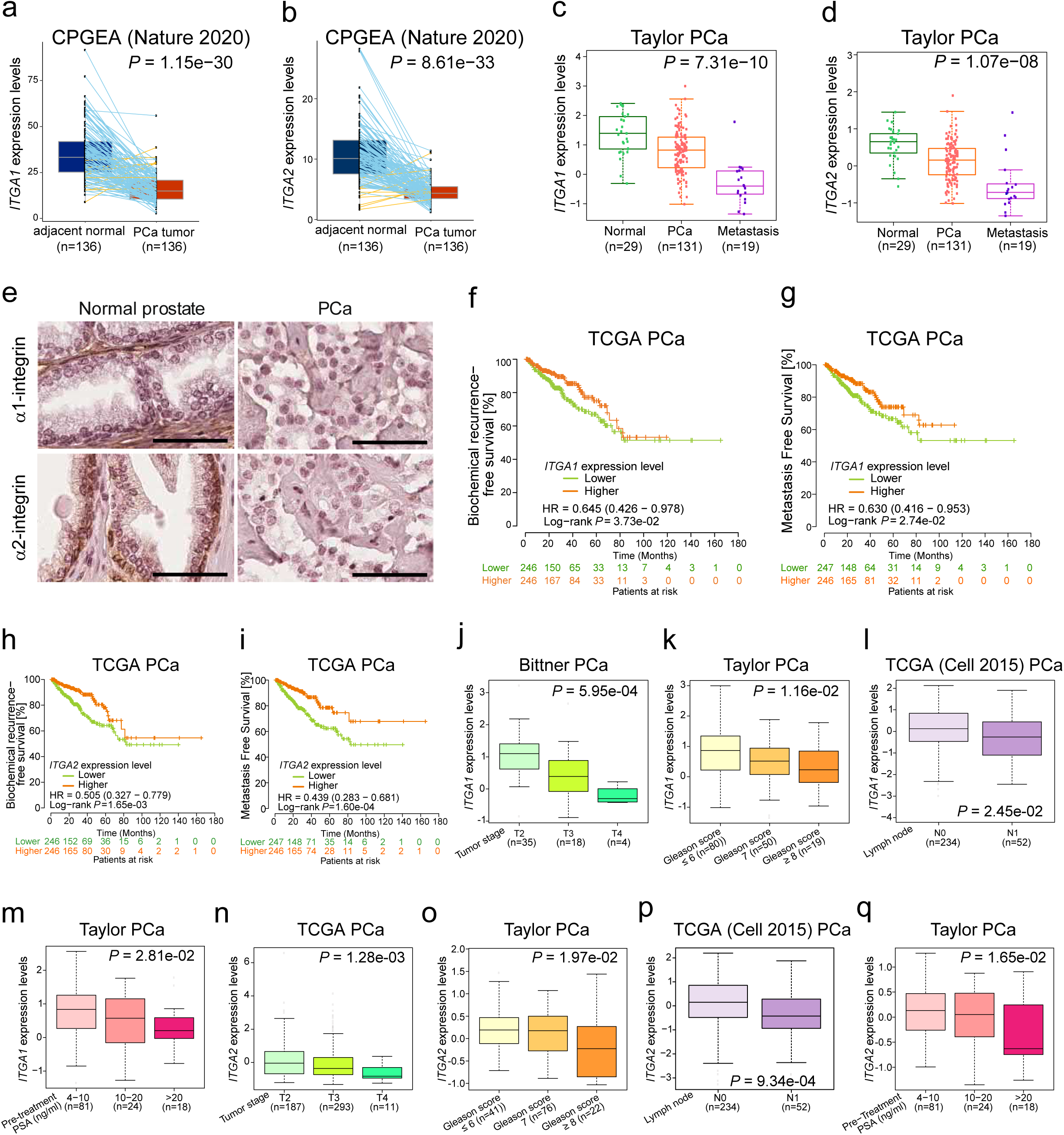
Downregulation of ITGA1 or ITGA2 correlates with PCa severity and progression. **(a-d)** Boxplots displaying expression levels of *ITGA1* (**a, c**) or *ITGA2* (**b, d**) downregulated during PCa development and progression. *P-*values were determined by the Kruskal-Wallis H test for comparing three or more groups (**a** and **b**) or the Mann-Whitney U test for comparing two groups (**c** and **d**). (**e**) Representative images of α1- and α2-integrin expression and localization in normal and PCa tissue. Scale bar=100µm. (**f-g**) Kaplan Meier plots indicate increased biochemical recurrence (**f**), and metastatic (**g**) risks for PCa patients with tumors expressing lower *ITGA1* levels. (**h-i**) PCa patients with decreased *ITGA2* expression level are associated with increased risks for biochemical recurrence (**h**), and metastasis (**i**). PCa patients were stratified into lower and higher expression groups by the median value of *ITGA1* or *ITGA2* expression levels. *P-*values were assessed by log-rank test. (**j**-**m**) Decreased *ITGA1* expression level correlates with higher tumor stage (**j**), Gleason score (**k**), lymph node metastasis (**l**) and PSA levels (**m**). (**n**-**q**) *ITGA2* downregulation is associated with higher tumor stage (**n**), Gleason score (**o**), lymph node metastasis (**p**) and PSA levels (**q**). *P* values were determined by Kruskal-Wallis H test.

### Loss of α1β1- and α2β1-integrins synergistically promote prostate epithelial cell invasive migration

Given the strong correlation of α1- and α2-integrin depletion and downregulation with PCa progression we analyzed the expression levels of α1- and α2-integrin levels in benign prostate epithelial cells (RWPE1) and the malignant PCa cell lines (DuCaP, PC3, VCaP, DU145 and 22Rv1) by western blotting (**Fig. 3a**). As expected, HPV18-immortalized benign RWPE1 cells expressed high levels of both α1- and α2-integrins. In contrast, brain metastasis-derived DuCaP and DU145 and prostate carcinoma cell line 22Rv1 displayed reduced levels of α1- and especially of α2-integrins. Two bone metastasis-derived cell lines showed reduced levels of either α1-(PC3) or α2-integrins (VCaP). In summary, cancer cell lines displayed downregulated levels of either one or both collagen-binding integrins.

**Figure 3.**
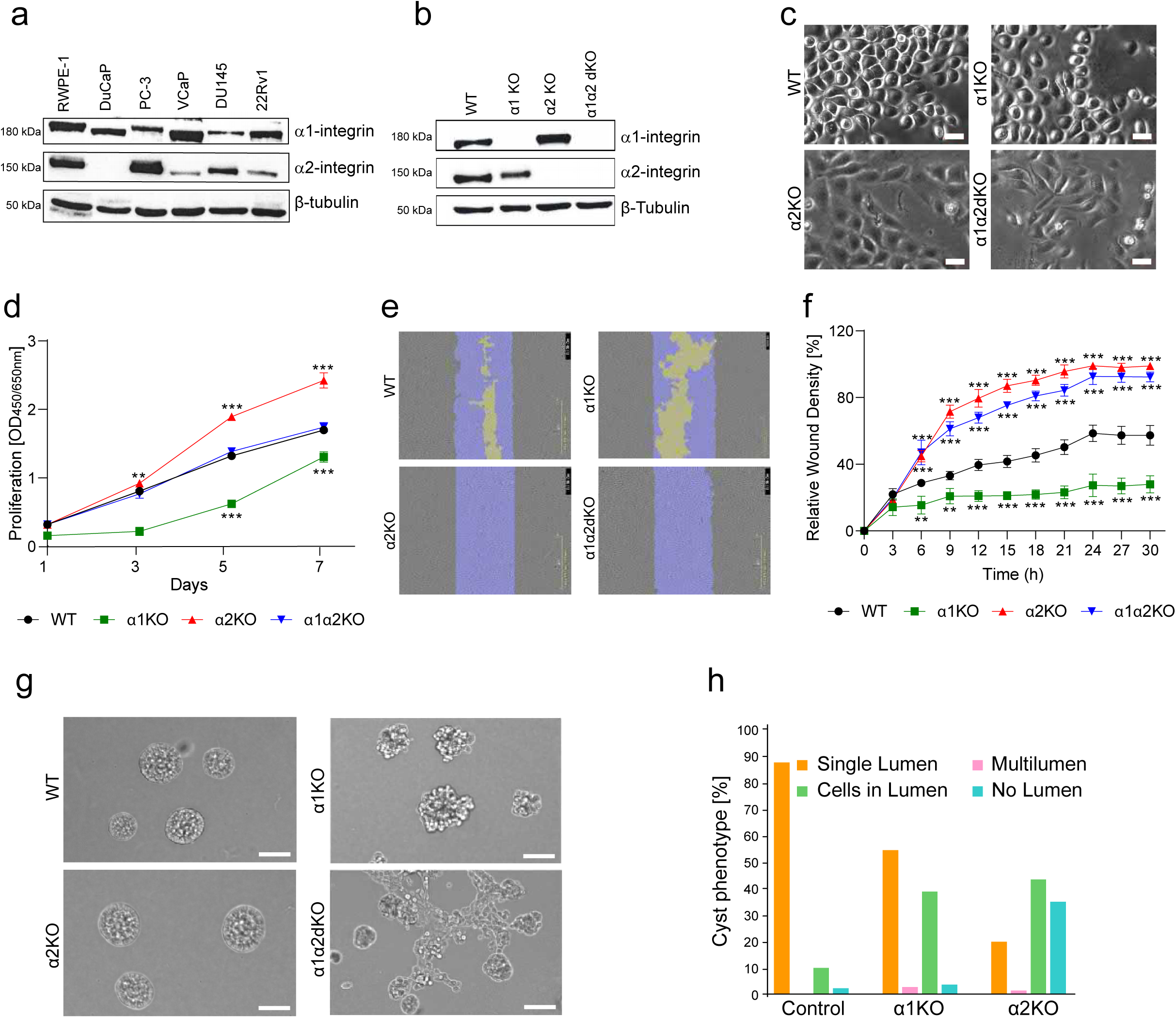
Loss of α1- or α2-integrins leads to different but synergistic phenotypes in prostate epithelial cells. (**a**) Level of α1- and α2-integrins in normal (RWPE1) and PCa (DuCaP, PC-3, VCaP, DU145, 22Rv1) epithelial cells. (**b**) RWPE1-WT, RWPE1-α1KO, RWPE1-α2KO and RWPE1α1α2dKO cell lysates were analyzed for the expression levels of α1- and α2-integrins. β-tubulin was used as a loading control. (**c**) RWPE1-WT, RWPE1-α1KO, RWPE1-α2KO and RWPE1-α1α2dKO cells grown on glass coverslips for 2 days were imaged using phase contrast microscopy. Scale bar=10µm. (**d**) Proliferation of the indicated RWPE1 cell lines were analyzed using an XTT assay. The data shows mean ±SD from three independent experiments performed in triplicates. (**e**) Migration of the different RWPE1 variants was analyzed using the IncuCyteS3 Scratch Wound module. (**f**) A plot showing the wound-closure dynamics of the indicated RWPE1 cell lines. The mean ± SD from a representative assay with 3 replicates is plotted in the graph. The assay was repeated thrice with similar results. (**g**) Phase contrast microscopy images of the indicated RWPE1 variants grown in 3D Matrigel for 7 days. Scale bar=50µm. (**h**) Quantitation of the 3D morphology analysis of RWPE1-WT, -α1KO and α2KO cell lines. RWPE1-α1α2dKO cells formed interconnected multicellular networks and could not be analyzed. Cysts were classified into 4 categories: cysts with a central hollow lumen, multilumen cysts, cysts with individual cells in the lumen and solid cysts with no visible lumen. The data shows the mean ± SD from three independent experiments in which at least 100 cysts per sample was analyzed. For all quantitative data in this figure Significance is indicated by asterisks: ∗ = p < 0.05; ∗∗ = p < 0.01; ∗∗∗ = p < 0.001.

To study the potential functional role of these integrins in prostate epithelial cells, we took benign RWPE1 cells, expressing high levels of both α1- and α2-integrins and generated α1-integrin knockout (α1KO) and α2KO RWPE1 cells using CRISPR/Cas9-mediated genome editing ^13^. Two independent gRNAs targeting different exons were designed for each gene and used as independent biological replicates in the subsequent experiments to confirm the specificity of the KO-phenotype and to exclude possible off-target effects. To address the potential redundance or synergistic functions of the two collagen receptors we also established a double knockout cell line (α1α2dKO). Specific loss of the target proteins was confirmed by Western blotting (**Fig. 3b** and **Fig. S4a**). In our culture conditions, parental RWPE1 cells and RWPE1 cells transduced with a control virus form a cobblestone-like cell monolayer upon confluency (**Fig. 3c**). Individual RWPE1-α1KO cells appeared somewhat rounder than control cells and while it took longer for RWPE1-α1KO cultures to reach confluency, they eventually formed a monolayer resembling that of the controls. RWPE1-α2KO cells, and especially the RWPE1-α1α2dKO cells, displayed an elongated shape reminiscent of mesenchymal or actively migrating cells (**Fig. 3c**). Curiously, RWPE1-α2KO cells reached confluency significantly faster than the other cell lines. To study cell proliferation in more detail, we performed an XTT assay. As expected, RWPE1-α2KO cells grew significantly faster and RWPE1-α1KO cells proliferated with slower kinetics when compared with control or α1α2dKO RWPE1 variants (**Fig. 3d**).

Given that both α1- and α2-integrins are collagen receptors, we examined their respective functional roles in collagen adhesion. RWPE1 control, -α1KO, -α2KO and -α1α2dKO cell variants were seeded onto collagen I-coated substrates and cell spreading in each sample was monitored. RWPE1 control, -α1KO and -α2KO cells all efficiently spread on collagen whereas the RWPE1-α1α2dKO cells lacking both α1- and α2-integrins failed to adhere and spread even after long (up to 12 hours) incubation periods (**Fig. S4c-d**). Thus, collagen-binding and cell spreading can be mediated by either α1- or α2-integrins but in the absence of both collagen adhesion is severely impaired in prostate epithelial cells. Next, we studied the migratory properties of the different RWPE1 variants using the IncuCyte S3 Scratch wound module. Confluent monolayers were scratched, and wound closure was monitored over a period of 30 hours. In line with the observed elongated morphology, wounds in the RWPE1-α2KO and RWPE1-α1α2dKO monolayers recovered significantly faster than in RWPE1 control monolayers (**Fig. 3e-f**). In contrast, RWPE1-α1KO cells showed significantly impaired wound closure capacity (**Fig. 3e-f**). Since α1α2dKO cells are incapable of spreading on collagen these results suggest that wound closure is mediated by cell migration on another cell secreted ECM, presumably fibronectin or laminin.

### Loss of α1β1- and α2β1-integrins leads to impaired acinar morphogenesis in 3D

PCa progression is associated with gradual loss of cell polarity leading aberrant glandular morphology and function. Benign RWPE1 cells form hollow cysts resembling acinar topology *in vivo* when cultured in 3-dimensional (3D) gels ^7, 52^. To assess morphology of control, α1-, α2- and α1α2-depleted RWPE1 cells, the different variants were seeded into 3D basement membrane extract (BME) gels and cultured for up to 7 days. The vast majority of control cells formed hollow cysts with smooth basal surface and an apical lumen occasionally containing few individual cells inside the lumen (**Fig. 3g-h** and **Fig. S4b**). Only half of the RWPE1-α1KO cells formed a central lumen while basal surfaces had a grape-like appearance possibly due to reduced cell-cell interactions **(Fig. 3g-h**). In contrast, 80% of RWPE1-α2KO cysts were smooth-surfaced solid clusters lacking visible apical lumens (**Fig. 3g-h**). Surprisingly, the 3D morphology of RWPE1-α1α2dKO cells could not be properly quantitated as they hardly formed any cysts and instead grew as solid cell clusters connected to each other via chains of collectively migrating cells (**Fig. 3g**). This phenotype suggests that α1/α2-dual depleted RWPE1 cells possess invasive properties in 3D cultures.

### Loss of α1β1- and α2β1-integrins promotes activation of the EMT pathway in prostate epithelial cells

To understand the role of *ITGA1*/*ITGA2* in PCa, we performed RNA-seq analysis of differentially expressed genes upon knocking out *ITGA1* or/and *ITGA2* with control in the RWPE1 cells (**Fig. 4a**). In each experimental group, two biological replicates were included and showed high correlations (**Fig. S5a-d**). The RNA-seq analysis identified 3073 upregulated and 2575 downregulated DEGs in RWPE1-α1α2dKO, 408 upregulated and 537 downregulated DEGs in RWPE1-α1KO and 2898 upregulated and 2317 downregulated DEGs in RWPE1-α2KO cells when compared with control (DESeq2, FDR < 0.05; **Fig. 4a**). To investigate potential functional categories of *ITGA1*/*ITGA2* KO target genes, we performed gene set enrichment analysis (GSEA) and revealed the epithelial-to-mesenchymal transition (EMT) pathway highly enriched in α1α2-dKO upregulated genes in the Hallmark gene sets (**Fig. 4b-c**). The interferon response pathways and EMT pathway were top enriched in the α1- or α2-KO upregulated genes, respectively (**Fig. S5e-f**). Given the strongest morphological phenotype and invasive behavior was observed in the RWPE1-α1α2dKO cells, we focused our analysis on these variants. To further consolidate these findings, we performed additional enrichment analysis in a multiple collections of annotated gene sets. The results constantly revealed the strong enrichment of genes upregulated upon α1α2-dKO for pathways relevant with cancer metastasis and EMT (**Fig. 4d**). Further expression profiling analysis of the EMT pathway revealed that the majority of genes in the EMT pathway were upregulated upon α1α2-dKO (**Fig. 4e**), which is in line with our above observations on the association of *ITGA1*/*ITGA2* copy number loss/del and downregulation with PCa tumor progression to advanced stages and metastasis (**Fig. 1f-m**; **Fig. 2**; **Fig. S1-3**). We next sought to evaluate whether the copy loss/del or expression levels of *ITGA1*/*ITGA2* directly correlates with PCa progression, and thus further performed correlation analysis with the EMT score that was reported to correlate with known EMT markers in PCa patient tumors (see **Methods**). The result showed that the EMT score was significantly upregulated in PCa patients with *ITGA1*/*ITGA2* copy loss/del (**Fig. 4f**). Consistently, *ITGA1*/*ITGA2* expression levels showed negative correlation with the EMT score (**Fig. 4g**). We further stratified PCa patients by the median expression value of *ITGA1*/*ITGA2,* and found that in line with previous findings, the EMT score was elevated in patient group with *ITGA1*/*ITGA2* low expression levels (**Fig. 4h**). To further validate these findings, we checked the expression levels of selected key EMT markers in the different cell lines by quantitative PCR. In line with the *in vitro* phenotype, RWPE1-α1α2dKO cells showed significant upregulation of *SLUG, TWIST, SNAII, CDH2, ITGA5* and *ITGAV* and downregulation of *CDH1* expression levels indicative of EMT (**Fig. 4i**). Of note, *CDH1* was strongly downregulated also in RWPE1-α1KO cells and RWPE1-α2KO cells whereas the RWPE1-α2KO cells showed curious downregulation of *SNAIL* and *CDH2* and neither of the single KOs displayed activation of the full EMT program relevant genes (**Fig. 4i**). Interestingly, the GSEA also revealed the TGFβ signaling pathways, representing a subset of the EMT pathway, significantly enriched in the upregulated genes identified by RNA-seq of α1α2dKO in the RWPE1 cells (**Fig. 4j-k**). Taken together, RNA-seq analysis highlighted activation of the EMT pathway and the associated TGFβ-signaling pathway as possible cellular transcriptional programs that underlie the potential higher tumorigenic potential of *ITGA1*/*ITGA2* loss & deletion in PCa patients and the invasive behavior in α1-/α2-integrin dual depleted prostate epithelial cells.

**Figure 4.**
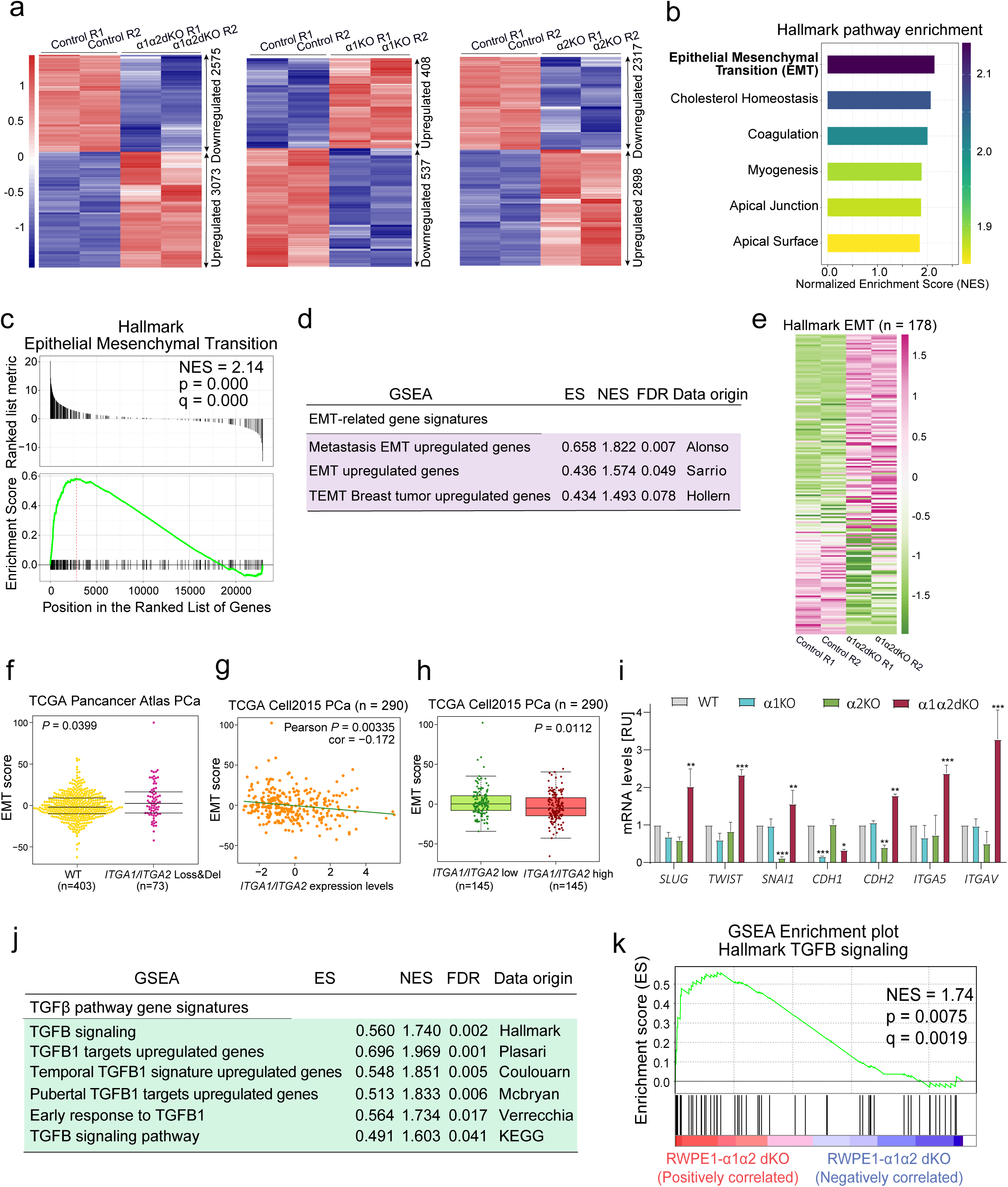
Loss of both α1- and α2-integrins activates EMT-associated pathways in prostate epithelial cells. (**a**) Heatmap representation of differentially expressed genes identified from the RNA-seq analysis in RWPE1-WT (control) and RWPE-α1α2dKO (left panel), -α1KO (middle panel) or -α2KO (right panel) cells. The false discovery rate (FDR) < 0.05. (**b**) Top-ranked pathways enriched in the upregulated genes form the GSEA analysis in RWPE1-a1a2dKO cells. Categories were ranked by the Normalized Enrichment Score (NES). (**c**) GSEA enrichment plot displaying the EMT pathway enriched in upregulated genes in RWPE1-a1a2dKO cells. (**d**) Validation of the metastasis/EMT-related pathway signatures in additional resources. (**e**) Heatmap representation of genes in the EMT pathways measured by RNA-seq in the RWPE1 cells with α1α2-dKO. (**f**) The EMT score is significantly elevated in PCa tumors with copy loss/deletion of both *ITGA1* and *ITGA2*. (**g**) The EMT score is negatively correlated with the expression levels of *ITGA1* and *ITGA2* in PCa. (**h**) The EMT score is downregulated in PCa tumors with higher expression levels of *ITGA1*/*ITGA2*. The EMT score and the *ITGA1*/*ITGA2* expression levels are calculated as the z-score sum. (**i**) Quantitative PCR analysis confirming regulation of the EMT pathway genes. The data is presented as mean ± SD of triplicate experiments. ∗ = p < 0.05; ∗∗ = p < 0.01; ∗∗∗ = p < 0.001. (**j**) The TGFβ signaling pathway was significantly enriched in the upregulated genes with α1α2dKO in the RWPE1 cells in multiple gene set resources. (**k**) GSEA enrichment plot demonstrating the TGFβ signaling pathway enriched in the upregulated genes in RWPE1 cells with α1α2dKO.

### RWPE1-α1α2dKO cells have elevated levels of TGFβ1 secretion leading to YAP1 activation

To directly assess activation of the TGFβ1 pathway in RWPE1-α1α2dKO cells we harvested culture medium from confluent cultures and analyzed levels of TGFβ1 from concentrated medium. TGFβ1 is secreted as a latent complex that can associate with several ECM proteins, such as fibronectin, and that needs to be mechanically or proteolytically activated prior to binding to TGFβ-receptor proteins at the cell surface ^53^. RWPE1-α1KO and RWPE1-α1α2dKO cells both secreted higher levels of TGFβ1 but only in the latter it was found in an activated form (**Fig. 5a-b** and **S6a-b**). We subsequently applied two independent approaches to experimentally confirm the role of TGFβ1 in invasive migration. First, we treated 3D cultures of control RWPE1 cells with recombinant TGFβ1 and found that this treatment induced similar invasive structures in control cells as were seen in RWPE1-α1α2dKO cells (**Fig. 5c**). Second, RWPE1-α1α2dKO cells grown in 3D BME gels were treated with TGFβ inhibitor, SB431542, that strongly inhibited the branching capacity of these cultures (**Fig. 5c**).

**Figure 5.**
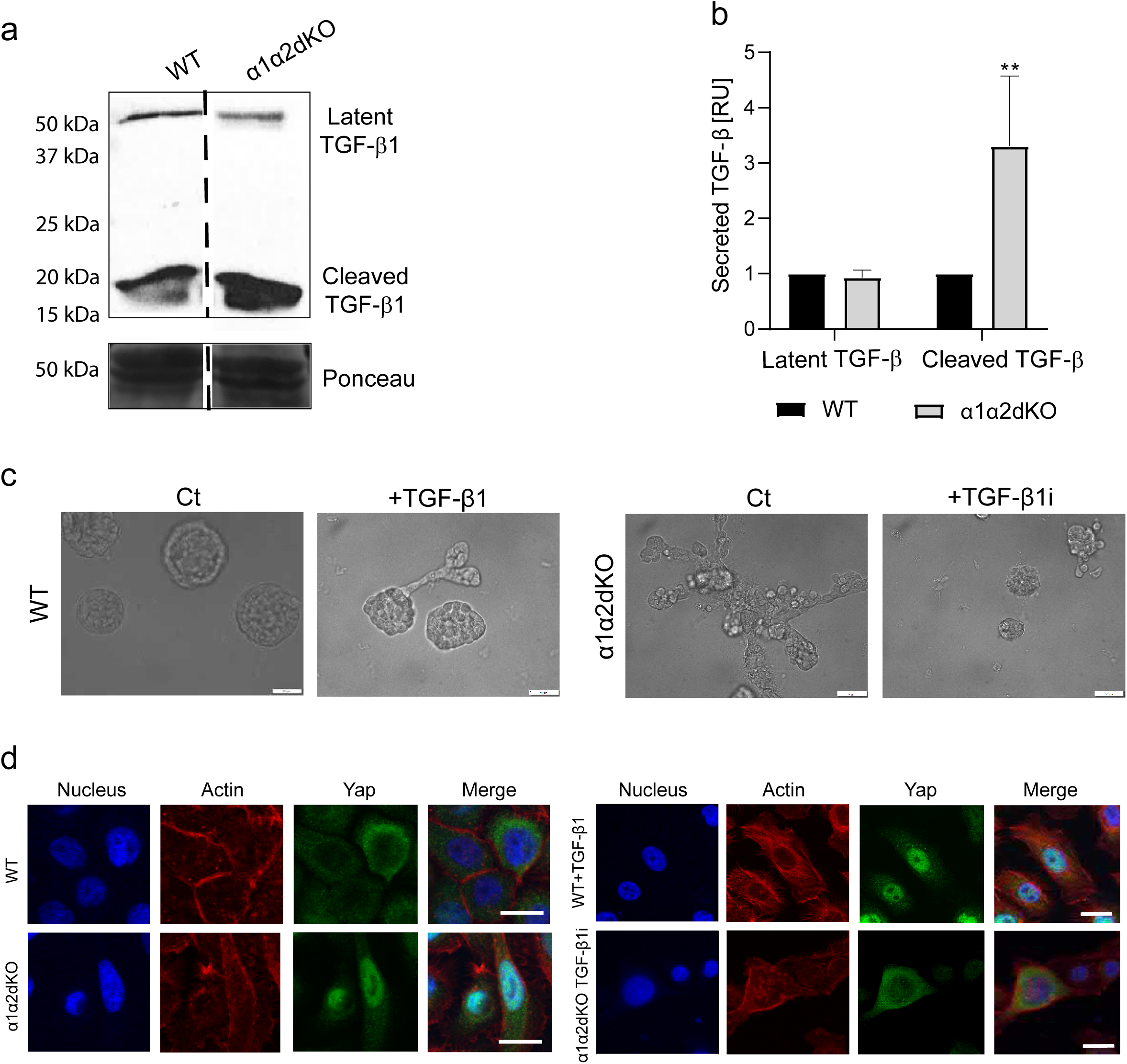
TGFβ activation drives invasive potential in α1- and α2-integrin-deficient prostate epithelial cells. (**a**) The culture medium from RWPE1-WT and RWPE1-α1α2dKO cells was harvested and concentrated as described in materials and methods. Secreted latent TGFβ and activated TGFβ was visualized by western blotting. The data is representative of three independent experiments with similar results. (**b**) Quantitative analysis of the secreted TGFβ. The data is presented as mean ± SD of triplicate experiments. ∗ = p < 0.05; ∗∗ = p < 0.01; ∗∗∗ = p < 0.001. (**c**) RWPE-1-WT cells were grown in 3D BME gel with or without 10ng of TGFβ1 for 7 days while RWPE1-α1α2dKO cells were grown in the presence or absence of 5µM TGFβ1 inhibitor (TGFβ1i). Scale bar: 50µm. (**d**) RWPE1-WT and RWPE1-α1α2dKO cells were grown on glass coverslips in the absence or presence of 10ng TGFβ1 or 5µM of TGFβ1i, respectively in coverslips for 2 days, fixed and stained for nuclei (blue), actin (red) and YAP1 (green). Scale bar:10µm.

The transcriptional coactivator YAP1 is a common oncogenic factor that promotes proliferation and migration in several cancer types including PCa ^54, 55^. Integrins are central regulators of YAP1 that is activated by the Hippo pathway and has been shown to crosstalk with the TGFβ pathway ^56–58^. Next, we analyzed YAP1 expression and subcellular localization in the different RWPE1 variants. In RWPE1 cells YAP1 is mainly cytoplasmic, while in α1α2dKO cells it was observed in the nucleus (**Fig. 5d**). To study if TGFβ activation regulates nuclear translocation of YAP1, we treated RWPE1 cells with TGFβ. This resulted in robust nuclear accumulation of YAP1 (**Fig. 5d**). In contrast, inhibition of TGFβ signaling prevented nuclear targeting of YAP1 in α1α2dKO RWPE1 cells (**Fig. 5d**). Thus, TGFβ activation was shown to drive nuclear translocation of YAP1. To investigate the functional role of YAP1, we generated lentiviral shRNA-vectors targeting YAP1 mRNA. Depletion of YAP1 levels in RWPE1 (**Fig. S7a,e** and α1α2-dKO RWPE1 cells (**Fig. S7f-h**) was confirmed at both mRNA and protein levels by qPCR and western blotting, respectively. Interestingly, YAP1-KD in RWPE1 and PC3 cells led to reduced expression levels of *ITGA1* and *ITGA2* (**Fig. S7d-e**). We next investigated the correlation between *ITGA1*/*ITGA2* and *YAP1* expression in the human prostate tumors and observed a significant linear positive expression correlation in multiple independent PCa cohorts (**Fig. S7k**). These findings suggest that YAP1 might contribute to the regulation of *ITGA1*/*ITGA2* transcription. Given that YAP1 has been reported to regulate transcription by interacting with TEAD-family members ^59^, we explored a large collection of genome-wide chromatin immunoprecipitation sequencing (ChIP-seq) data derived from PCa and other tissues including breast and lung to study the regulatory mechanisms at the *ITGA1*/*ITGA2* loci. We found an enrichment of chromatin binding of TEAD1 and YAP1 transcription factors in the proximity of *ITGA1* and *ITGA2* promoter regions (**Fig. S7l**). To study whether YAP1 is required for the activation of TGFβ signaling in α1α2dKO cells, we generated YAP1-depleted α1α2dKO RWPE1 cells (α1α2dKO-YAP1KD). While α1α2dKO-YAP1KD cells showed reduced YAP1 levels, no significant effect was seen on their ability for secreting and activating TGFβ1 (**Fig. S7i**). However, downregulation of YAP1 did lead to reduced proliferation of α1α2dKO cells suggesting that YAP1 activation contributes to the proliferative capacity of α1- and α2-integrin depleted PCa cells (**Fig. S7j**).

### α1α2KO cells have enhanced metastatic capacity in vivo

The parental RWPE1 cells are benign and do not form tumors *in vivo* ^60^. Given the apparent invasive properties of α1- and α2-depleted RWPE1 cells *in vitro*, we next assessed the metastatic capacity of control, α1KO, α2KO and α1α2dKO RWPE1 cells using the tail-vein injection model in immunocompromised SCID mice. All the cell lines were transduced to express luciferase and GFP to allow *in vivo* visualization. After 4 weeks the mice were sacrificed, and the lungs were imaged using IVIS. Significant tumor foci were seen only for the α1α2dKO-injected group and weak signals were observed in some of the lungs in the α2KO group (**Fig. 6a**). Interestingly, detailed histological analysis of the lungs revealed several micrometastases in all but one mouse of the α1α2dKO group and in two α2KO samples. A single very small lesion was found in one to the α1KO samples (**Fig. 6b**). As expected, no lesions were found in the RWPE1 control samples. The sizes of micrometastases in α1α2dKO samples were significantly larger than those found in α2KO cells (**Fig. 6c**). Given that the capacity of tumor cells to survive in circulation is crucial for their metastatic capacity, we next analyzed the number of circulating tumor cells (CTCs) and thus collected the blood from each mouse and quantitated the number of CTCs. Blood could not be collected from one mouse injected with α1α2dKO cells. This analysis revealed that significantly more CTCs were detected in α1α2dKO and α2KO groups (**Fig. 6d**). Collectively, these results provide supporting evidence that the benign prostatic epithelial cells with co-deletion of α1 and α2 integrins showed enhanced metastatic potential via *in vivo* tumorigenesis studies in mice.

**Figure 6.**
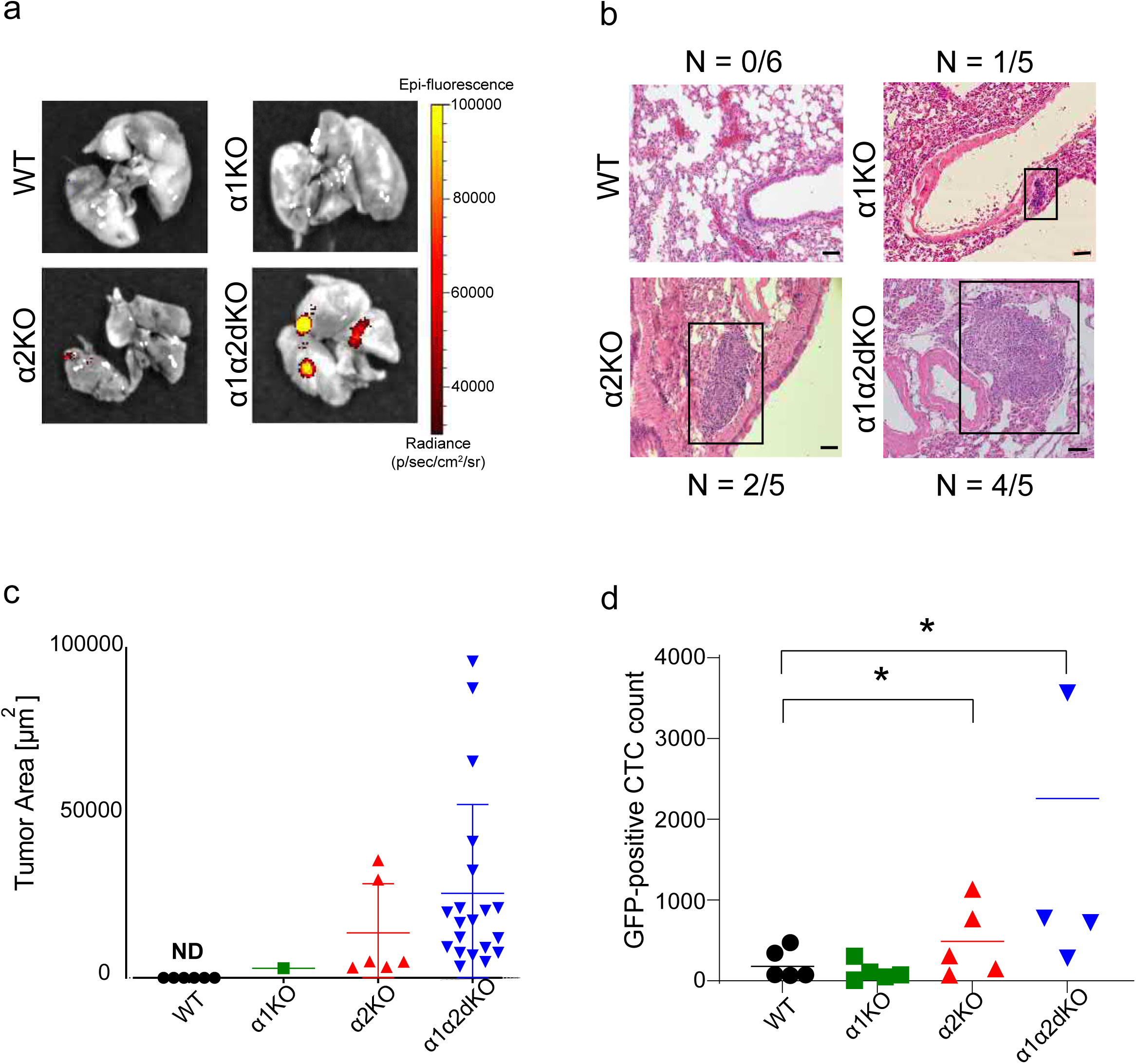
Deletion of α1- and α2-integrins in benign RWPE1 prostate epithelial cell line promotes formation of lung micro-metastases. (**a**) 1x10^5^ RWPE1-WT, -α1KO, -α2KO and - α1α2dKO cells expressing GFP and luciferase were injected into the tail veins of 6 immunodeficient SCID-mice per group. After four weeks, mice were sacrificed and the luciferase signals originating from cancer cells were measured using IVIS imaging system. (**b**) HE-stained lungs sections were analyzed for presence of metastatic lesions. Scale bar is 50µm. (**c**) The area covered by individual lung metastatic lesions were determined for each cell variant from HE-stained lung sections as described in experimental procedures. The data shows mean ± SD. (**d**) FACS-analysis of the number of circulating GFP-positive RWPE-1 cell variants recovered from the blood of sacrificed SCID-mice. For RWPE1-α1α2dKO cells, blood samples were obtained from only 4 mice. Data shows mean ± SD. ∗ = p < 0.05; ∗∗ = p < 0.01; ∗∗∗ = p < 0.001.

### TEAD1 transcriptionally correlates with ITGA1 and ITGA2 and directly regulates their expression via chromatin binding

To get insight into transcriptional regulation of *ITGA1* and *ITGA2*, we performed a genome-wide co-expression analysis of *ITGA1*/*ITGA2* in multiple PCa datasets to find out potential transcription factors that regulate *ITGA1*/*ITGA2*. Interestingly, TEAD1 emerged as the top positively co-expressed transcription factor gene with *ITGA1*/*ITGA2* in TCGA cohort consisting of 498 localized PCa tumors (**Fig. 7a-e**). Consistently, this finding was verified in another PCa cohort comprised of 150 tumor samples (**Fig. S8a-e**). In contrast, we observed that the expression levels of the other members of the *TEAD* gene family, *TEAD2-4*, were not correlated with *ITGA1* and *ITGA2* genes (**Fig. 7a-b** and **Fig. S8a-b**). This finding suggested that TEAD1 as the Hippo pathway transcription factor and YAP1 interacting protein, could be a key transcriptional driver of *ITGA1* and *ITGA2*.

**Figure 7.**
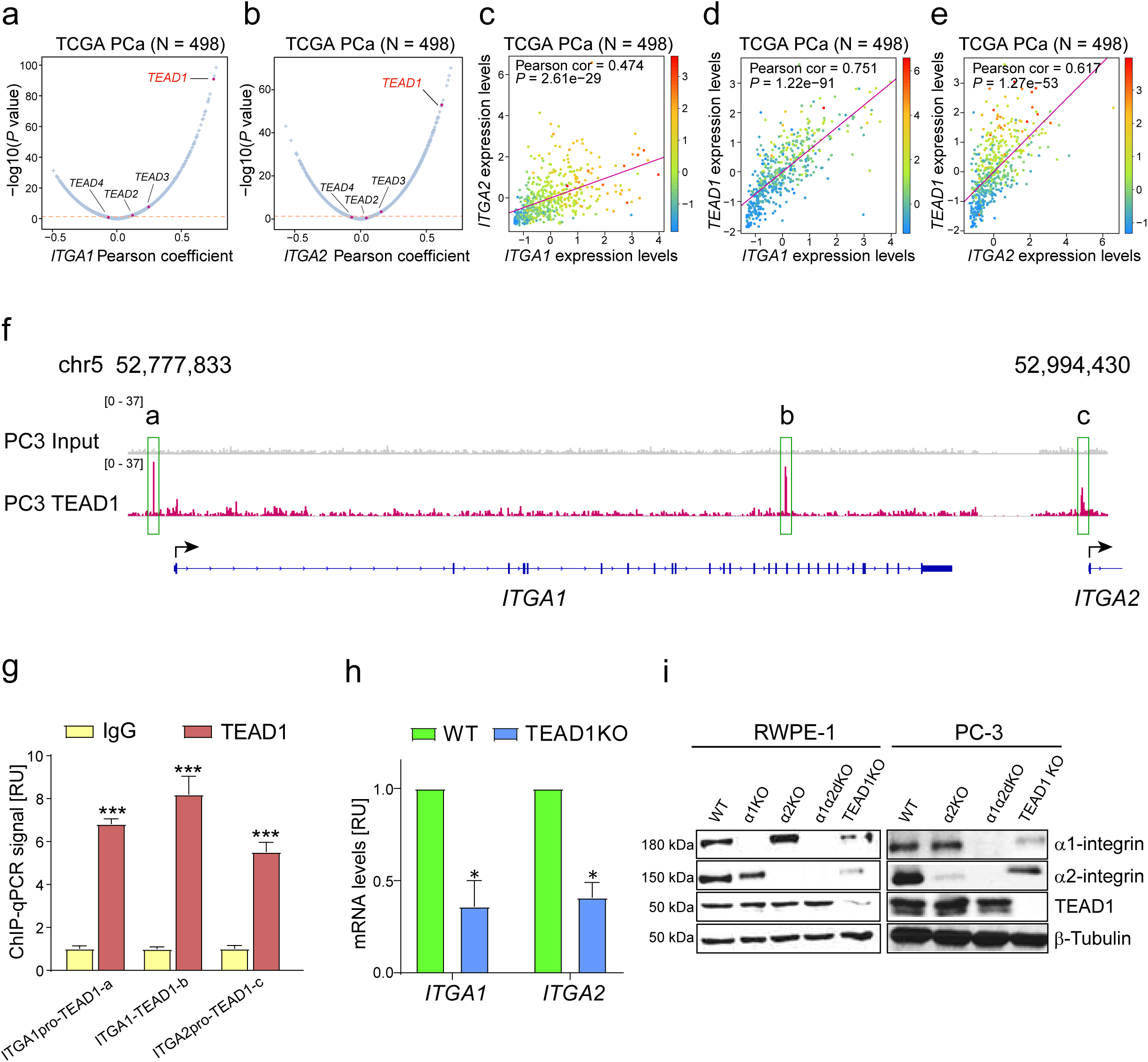
TEAD1 regulates *ITGA1* and *ITGA2* expression. (**a**-**b**) A genome-wide co-expression analysis identifies strong positive expression correlation of *ITGA1*(**a**) *or ITGA2*(**b**) with *TEAD1,* but not with *TEAD2, TEAD3* or *TEAD4* in the TCGA cohort. The X-axis demonstrates Pearson coefficient while Y-axis represents –log10 (*P* value). (**c-e**) Expression correlation analysis revealed significant positive correlation between *ITGA1* and *ITGA2* (**c**)*, ITGA1* and *TEAD1*(**d**), and *ITGA2* and *TEAD1* (**e**) in the TCGA data set. The color bars on the right side of figures indicate the expression level of *TEAD1*, *ITGA2* and *ITGA1*, respectively. *P*-values assessed by the Pearson’s product-moment correlation test. (**f**) Genome browser representation of ChIP-seq signals of transcription factor TEAD1 enriched in the promoters and surrounding regulatory regions of *ITGA1* and *ITGA2*. (**g**) ChIP-qPCR validation of TEAD1 at the three binding sites. (**h**) Knockdown of TEAD1 in the RWPE1 cell line downregulates expression of *ITGA1* and *ITGA2*. The data is presented as mean ± SD of triplicate experiments each with duplicates. (**i**) Western blotting of α1- and α2-integrins and TEAD1 in the indicated RWPE1 and PC3 cell variants. The blot is representative of three independent experiments with similar results. ∗ = p < 0.05; ∗∗ = p < 0.01; ∗∗∗ = p < 0.001.

To experimentally prove this, we performed TEAD1 ChIP-seq in the PC-3 cell line as previously described ^61, 62^. Motif analysis showed top-enriched DNA-binding motif of the TEAD family transcription factors (**Fig. S8f**), demonstrating the success and reliability of the ChIP-seq experiment. The subsequent genomic annotation analysis of ChIP-seq peaks illustrated that TEAD1 chromatin occupancies were greatly enriched in the gene proximal promoter regions compared to the genome background (**Fig. S8g**). Notably, among TEAD1 chromatin binding regions, we observed an enrichment of TEAD1 binding sites at the bi-directional promoter of *ITGA1* and *ITGA2* genes and within the ITGA1 gene body (**Fig. 7f**). To further validate the binding of TEAD1 at *ITGA1* and *ITGA2* gene regulatory regions, we performed ChIP-qPCR analysis in PC-3 cells as described previously (Huang et al., 2014). This analysis confirmed TEAD1 chromatin binding at the three regions determined by ChIP-seq (**Fig. 7g**). To further evaluate TEAD1-mediated regulation of *ITGA1/ITGA2* expression, we established TEAD1-KO cells of RWPE1 and PC3. Strong reduction of *ITGA1* and *ITGA2* mRNA expression was observed in TEAD1-KO RWPE-1 cells (**Fig. 7h**). Furthermore, western blot analysis confirmed robust downregulation of α1- and α2-integrins in RWPE1- and PC3 TEAD1-KO cells (**Fig. 7i**). Collectively, these results demonstrate that TEAD1 directly regulates the expression of *ITGA1/ITGA2* and this gene expression control might also hold true in the clinical setting due to their robust high co-expression rates across multiple independent cohorts of PCa.

### TEAD1 expression correlates with ITGA1 and ITGA2 expression and is lost during PCa progression

The expression levels of *TEAD1* and *ITGA1*/*ITGA2* were found to be strongly correlated and we experimentally confirmed transcriptional control of *ITGA1*/*ITGA2* expression by TEAD1. Given that ITGA1/ITGA2 copy loss and downregulation in turn correlated with PCa severity, we next studied whether *TEAD1* genomic alterations and expression levels directly correlate with PCa clinical characteristics. Intriguingly, similar to ITGA1/ITGA2, we found that *TEAD1* also indicated highly frequent genomic loss/deletion and its expression levels displayed linearly positive correlation with the genomic alterations (**Fig. 8a** and **Fig. S9a-b**). Notably, PCa patients with *TEAD1* copy loss/del showed significantly higher risk for biochemical recurrence (**Fig. 8b**). In line with the association of *TEAD1* copy loss/del with PCa clinical severity, we observed that *TEAD1* expression levels were gradually decreased upon PCa development and progression compared to normal prostates (**Fig. 8c-d** and **Fig. S9c-e**). Moreover, we also observed that *TEAD1* expression levels were significantly decreased with PCa severity including more advanced tumor stage (**Fig. 8e)**, higher Gleason score (**Fig. 8f)**, lymph nodes (**Fig. 8g)** and PSA levels (**Fig. 8h)**. Given that *TEAD1* was downregulated upon disease progression, we next thought whether *TEAD1* expression may possess prognostic value. We thereby conducted the Kaplan-Meier survival analysis and found that low *TEAD1* expression significantly correlated with an increased risk of biochemical recurrence (**Fig. 8i-j** and **Fig. S9f**) and metastasis (**Fig. 8k)** in PCa patients. Next, we utilized our TMA to investigate TEAD1 expression in the prostate and the potential correlation between TEAD1 levels and PCa progression. In benign samples TEAD1 was highly expressed in the nucleus of basal epithelial cells (**Fig. 8l**). Some nuclear TEAD1 expression was also seen in stromal cells (**Fig. 8l** and **Fig. S3a**, **d**). In contrast, we observed little or no TEAD1 staining in cancer tissues (**Fig. 8l** and **Fig. S3a**, **d**). In line with these observations, western blot analysis revealed high levels of TEAD1 in the benign RWPE1 cells whereas most malignant PCa cell lines exhibited lower Tef-1 levels (**Fig. 8m**). Curiously, TEAD1 levels in the different PCa cell lines appeared to correlate with α2-integrin levels (**Fig. 3a**).

**Figure 8.**
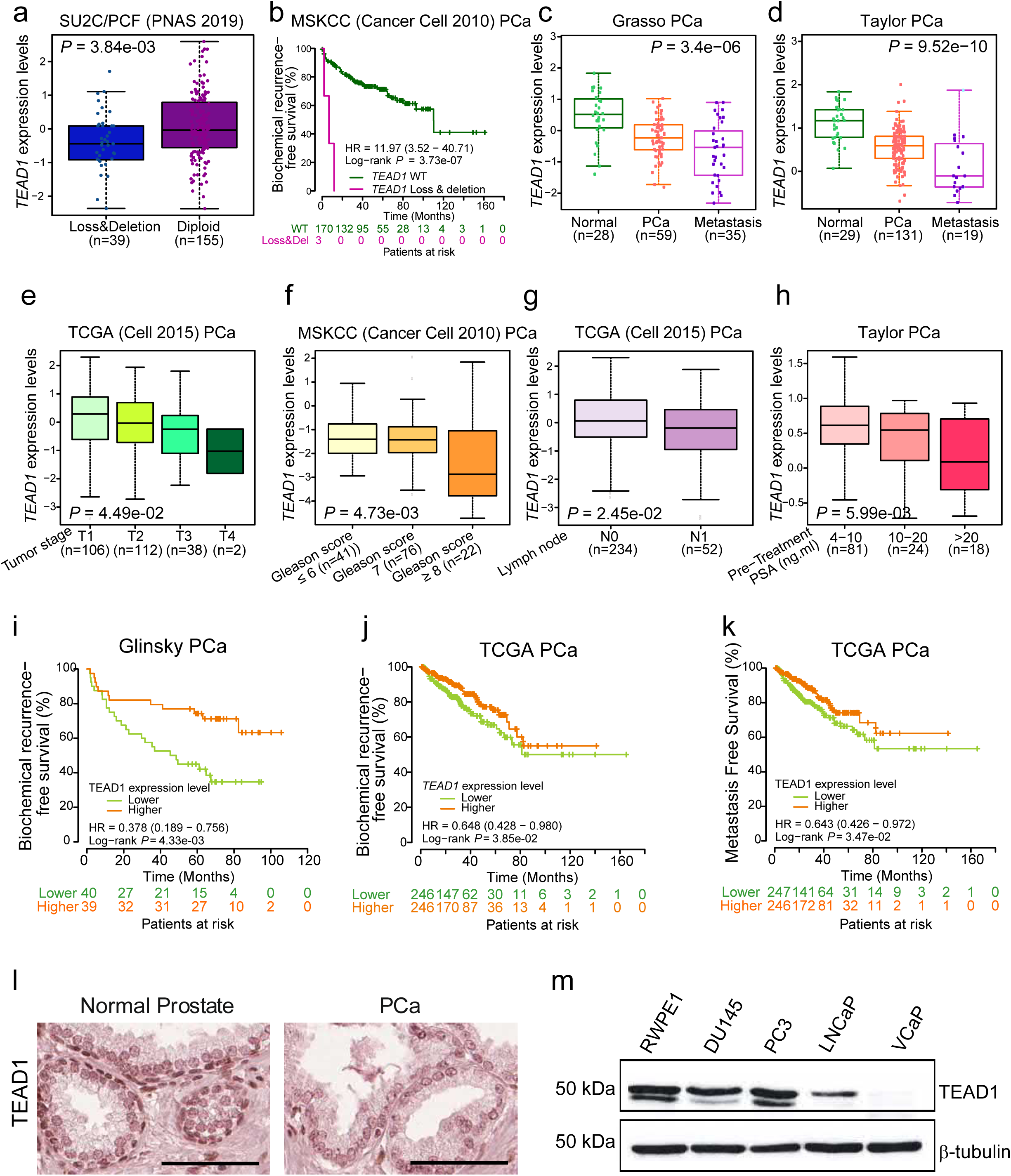
*TEAD1* is highly co-expressed with *ITGA1* and *ITGA2* and is downregulated during PCa development and progression. **(a)** *TEAD1* expression level are downregulated upon copy loss/del in PCa. *P* value was examined by the Mann-Whitney U test. (**b**) PCa patients with *TEAD1* copy number loss/deletion show higher risks for biochemical recurrence. *P*-value was assessed by the log-rank test. (**c**-**d**) *TEAD1* expression levels are significantly decreased upon PCa tumor progression to metastasis. (**e-h**) *TEAD1* downregulation correlates with PCa tumor progression to high tumor stages (**e**), Gleason score (**f**), lymph node metastasis (**g**) and PSA levels (**h**). *P* values were determined by Kruskal-Wallis H test. (**i-k**) Lower *TEAD1* expression levels in PCa patients are associated with higher risks for biochemical recurrence (**i**-**j**) and metastasis (**k**). *P*-values were assessed by the log-rank test. (**l**) Representative images of the TEAD1 protein expression in normal and PCa cancer tissue. Scale bar=100µm. (**m**) Western blot analysis of the TEAD1 expression in a benign (RWPE-1) and malignant (DU145, PC3, LnCap, VCap) epithelial prostate cell lines. The blot is representative of three independent experiments with similar results.

### Loss of TEAD1 phenocopies the α1α2KO cells

Given the strong correlation of *TEAD1* and *ITGA1*/*ITGA2* expression levels and the observed regulatory circuit of *TEAD1*-*ITGA1*/*ITGA2* we next examined whether the RWPE1-TEAD1-KO cells display similar transformed phenotypes observed in RWPE1-α1α2dKO cells. While not as striking as seen in α1α2dKO cells, confluent RWPE1-TEAD1-KO cells had slightly more elongated shape (**Fig. 9a**). Like α1α2-dKO cells, TEAD1-KOs showed similar proliferative capacity when compared with the controls (**Fig. 9b**) but migrated significantly faster than the controls in the scratch wound assay (**Fig. 9c**). Remarkably, 3D BME-grown RWPE1-TEAD1-KO cells displayed branching morphogenesis suggesting an invasive phenotype (**Fig. 9d**). The branching of TEAD1-KO RWPE1 cells was shown to be TGF-β1-dependent as TGF-β1i completely blocked invasive migration in 3D (**Fig. 9e**). TEAD1-KO RWPE1 cells also displayed autocrine secretion and activation of TGFβ1 (**Fig. 9f**). The tumorigenicity of RWPE1-KO cells was analyzed using the tail vein injection model as described for the integrin mutants above. RWPE1-TEAD1-KO cells formed micrometastases in all the injected mice and the size of these lesions was similar as was seen for α1α2dKO lesions (**Fig. 9g-h**). Moreover, CTCs were recovered from the blood of every mouse injected with TEAD1-KO RWPE1 cells (**Fig. 9i**).

**Figure 9.**
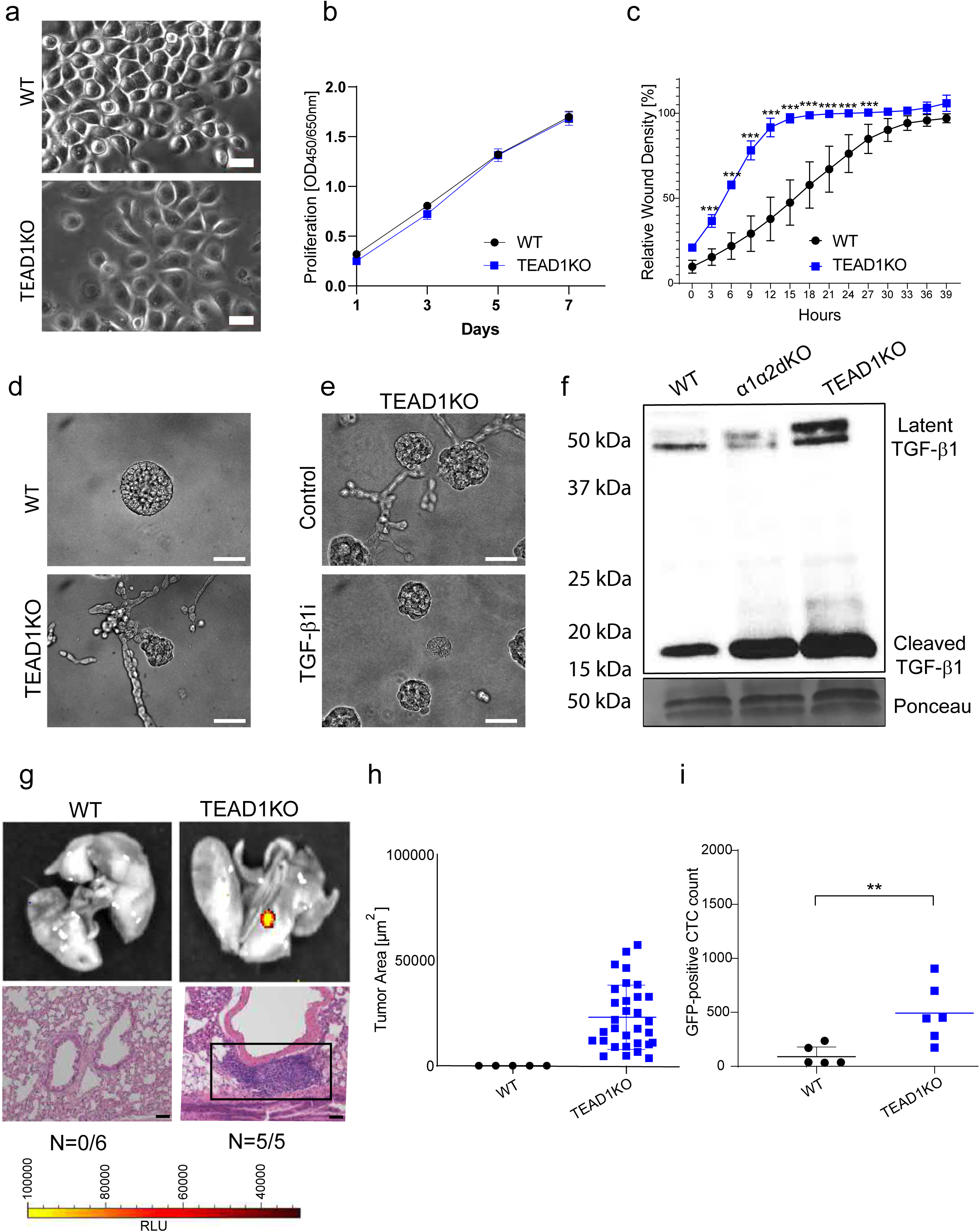
Loss of *TEAD1* phenocopies the dual loss of α1- and α2-integrins *in vitro* and *in vivo*. (**a**) RWPE1-WT and RWPE1-TEAD1KO cells grown on glass coverslips for 2 days were imaged using phase contrast microscopy. Scale bar=10µm. (**b**) Cell proliferation analysis of RWPE1 and RWPE1-TEAD1KO cells was done using XTT assay. The data shows mean ±SD from three independent experiments performed in triplicates. (**c**) Cell migration analysis using the IncuCyte S3 scratch wound module. The plot shows the mean ± SD from a representative assay done in triplicate. The assay was done three times with similar results. (**d**) Phase contrast microscopy images of RWPE1-WT and RWPE1-TEAD1KO cells grown in 3D BME gels for 7 days. The scale bar is 50µm. (**e)** RWPE1-TEAD1KO cells were grown in the presence or absence of 5µM TGFβ1 inhibitor (TGFβ1i) as described in (d). Scale bar is 50µm. (**f**) Culture medium was harvested and concentrated from RWPE1-WT, RWPE1-α1α2dKO and RWPE1-TEAD1KO cells were (**g**) Luciferase expressing RWPE1-WT and -TEAD1KO cells were imaged using the IVIS imaging system. Scale bar: 50µm. (**h**) The area covered by individual lung metastatic lesions was determined for each cell variant from HE-stained lung sections as described in experimental procedures. The data shows mean ± SD. (**i**) FACS analysis of the circulating WT and TEAD1-KO RWPE1 cells collected from the blood of sacrificed SCID mice. Total GFP-positive cell count from the sample of each mouse and the mean for each group is shown in the plot. Data shows mean ± SD. ∗ = p < 0.05; ∗∗ = p < 0.01; ∗∗∗ = p < 0.001.

### Synergistic effects of *ITGA1*, *ITGA2* and *TEAD1* downregulation promotes PCa progression and severity

We described above that downregulation of *ITGA1*, *ITGA2* or *TEAD1* demonstrated apparent associations with poor prognosis of PCa patient (**Fig. 2f-i**, **Fig. S2i-n**, **Fig. 8i-k** and **Fig. S9f**), and thus sought to explore whether *ITGA1*, *ITGA2* or *TEAD1* expression levels may hold predictive value for stratifying PCa patients with low- and high-risks. We hence stratified several cohorts of PCa patients according to the Gleason scores and investigated the potential correlation between gene expression and patient prognosis. The result suggested an explicit predictive value of *ITGA1*, *ITGA2* or *TEAD1* mRNA levels for patient prognosis in PCa group with a Gleason score of 7 (intermediate risk, **Fig. 10a-f**), but not for the low-risk cases with Gleason score ≤6 (**Fig. S10a-f**) or high-risk group with Gleason score ≥8 (**Fig. S10g-l**). These results indicate *ITGA1*, *ITGA2* or *TEAD1* as a potential independent prognostic marker in distinguishing PCa patients who are classified with the intermediate risks, representing the most difficult patient group in clinic to avoid the overtreatment.

**Figure 10.**
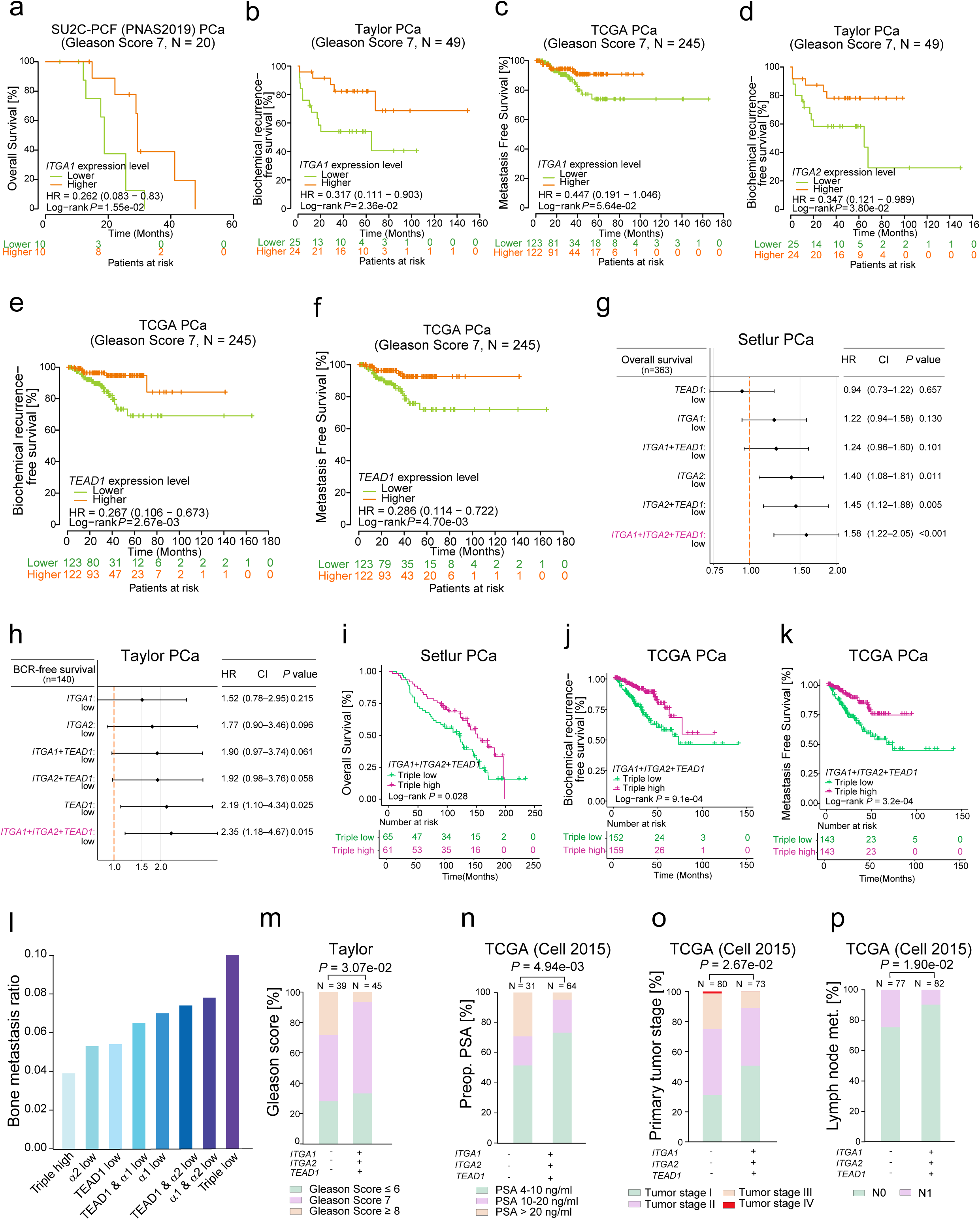
The prognostic value of *ITGA1*, *ITGA2* and *TEAD1* for PCa patient risk stratification and their synergistic impact on PCa in the clinical settings. **(a-c)** Lower expression level of *ITGA1* exhibit predictive values for overall survival (a), biochemical recurrence (b), and metastasis (c) in PCa patient group with an intermediate risk (Gleason Score 7). (d) PCa tumors with lower expression level of *ITGA2* is associated with decreased biochemical recurrence-free survival in group with Gleason score 7. (e-f) Lower expression levels of *TEAD1* holds predictive values for increased risks of biochemical recurrence (e) and metastasis (f) in PCa patients with an intermediate risk (Gleason score 7). *P*-values were assessed by log-rank tests. (g-h) Forest plots demonstrating triple-low expression of *ITGA1*, *ITGA2* and *TEAD1* with high hazard ratio for the overall survival (g) and biochemical recurrence-free survival (h) of PCa patients. (i–k) PCa patients with triple-low expression of *ITGA1*, *ITGA2* and *TEAD1* are associated with shorter overall survival (i), increased risks of biochemical recurrence (j) and metastasis (k). (l) Bone metastasis ratio for patients stratified based on high or low α1-integrin, α2-integrin and TEAD1 expression status. (m–p) PCa patients with triple-low expression of *ITGA1*, *ITGA2* and *TEAD1* are associated with advanced tumors including Gleason Score (m), PSA levels (n), tumor stage (o), and lymph node metastasis (p).

Although we found that reduced expression of *ITGA1*, *ITGA2* or *TEAD1* correlated with PCa severity, it remains unclear whether loss of these three genes have synergistic effects on tumor severity compared with loss of each individual gene. Therefore, we calculated the expression sum of the three genes in pairwise or triple-wise manner and examined the effect on patient prognosis. Intriguingly, the results showed that triple low expression of the three genes demonstrated highest hazard ratio for the overall survival (**Fig. 10g**) and biochemical recurrence (**Fig. 10h**) of PCa patients. To solidify our findings, we stratified PCa patients with triple low or -high expression levels of *ITGA1*, *ITGA2* and *TEAD1* and conducted the Kaplan–Meier analysis in multiple PCa cohorts. We found that patients with simultaneous triple low expression of *ITGA1*, *ITGA2* and *TEAD1* demonstrated significantly shorter overall survival (**Fig. 10i**), higher risk of biochemical recurrence (**Fig. 10j** and **Fig. S10m**), and metastasis (**Fig. 10k**). In addition, upon stratification of our inhouse TMA patients with expression data of all the three proteins combined, we found that patients with triple negative/low expression have an increased risk to develop bone metastasis when compared with patients with a loss of expression of only one or two of the markers. The biggest difference was found when the triple-negative/low patient groups were compared with triple-positive patient group (**Fig. 10l**). To further explore the synergistic effects of the loss of the three genes in clinical settings, we examined the correlation with several clinical features including Gleason score, PSA, tumor stage, and lymph nodes in PCa cases. The results constantly showed that PCa patients with triple low expression of *ITGA1*, *ITGA2* and *TEAD1* had higher Gleason score (**Fig. 10m**), PSA levels (**Fig. 10n** and **Fig. S10n**), more advanced tumor stages (**Fig. 10o**) and lymph node metastases (**Fig. 10p**). Taken together, these findings suggested that identification of the *ITGA1*/*ITGA2*/*TEAD1*-triple negative PCa cases may have potential to detect aggressive forms, especially in the most problematic Gleason score 7 PCa cases with uncertain PCa pathogenesis. PCa patient group with triple low expression of the *ITGA1/ITGA2*/*TEAD1*-axis was markedly associated with poor prognosis, increased disease severity and PSA levels, a biomarker for early detection of PCa. Despite its usefulness, PSA alone is not a reliable diagnostic tool ^63^. Our data suggests that the analysis of *ITGA1/ITGA2*/*TEAD1* expression in the PCa tissue could be considered as novel additional biomarkers helping to make more accurate treatment decisions in the clinic for PCa patients with high PSA levels and intermediate Gleason score 7.

## DISCUSSION

PCa progression involves drastic changes in cell-ECM contacts. A key feature observed in PCa is the loss of hemidesmosomes mediated by α6β4-integrins ^7, 64–66^. The meta-analysis performed here revealed a general downregulation of the integrin signaling pathway. While it is thought that invasive PCa cells need to interact with the stromal ECM rich in collagen, we found that the α-subunits of the main collagen-binding integrins, α1 and α2, were also strongly downregulated. Importantly, we demonstrated that loss of both α1- and α2-integrins not only correlated with PCa progression but also contributed to tumorigenesis by inducing autocrine secretion and activation of TGFβ leading to EMT. *ITGA1* and *ITGA2* are neighboring genes in the integrin collagen receptor locus on human chromosome 5q11.2. Certain single nucleotide polymorphisms (SNPs) within this region have been found to be associated with increased PCa risk ^67^. We report here that deletion of this locus is a relatively common (>2%) event in prostate, ovarian and esophageal cancers and deletion and loss as well as downregulation of both *ITGA1* and *ITGA2* correlate with development of an aggressive form of PCa.

Here we identified Tef-1, encoded by *TEAD1* gene, as the key regulator of *ITGA1*/*ITGA2* transcription. Out of the four different *TEAD* family members, only *TEAD1* expression was strongly correlated with *ITGA1* and *ITGA2* expression. All the four TEAD proteins have been implicated in coordination of especially the Hippo signaling pathway, but TEADs regulate also Wnt-, TGFβ- and epidermal growth factor receptor (EGFR) signaling pathways. Surprisingly, while TEAD overexpression has frequently been linked to tumorigenesis, we observed here that *TEAD1* downregulation or deletion was strongly associated with aggressive PCa. Our data suggest that deletion or downregulation of *TEAD1* contributes to PCa progression by abrogating α1- and α2-integrin expression. In contrast to our data, a previous study reported that upregulation of *TEAD1* correlated with worse PCa prognosis ^68^. The reasons for these discrepant results are unclear but the specificity of the TEAD1/Tef-1 antibodies used for immunohistology could perhaps be one contributing factor. In addition to histological studies, our current study included analyses of *TEAD1* mRNA levels and genomic deletion or loss of *TEAD1* with all the data supporting a tumor suppressive function for *TEAD1* in PCa. Curiously, in agreement with our data, Knight *et al*. also reported aberrant 3D morphogenesis in TEAD1-depleted RWPE1 cells. Thus, it seems clear that the role of TEADs in PCa need further scrutiny. It is also noteworthy, that despite early promise, recent studies addressing the pharmacological potential of targeting the TEAD/YAP1-axis have raised concerns related to activation of alternative oncogenic pathways as seen for *TEAD1*-KO cells in our study ^69^.

While *TEAD1* loss mediates some of its tumorigenic effects by influencing α1- and α2-integrin levels there were also significant differences in the mechanisms. First, we found that α1α2dKO led to TGFβ-driven EMT and nuclear translocation of YAP1 to promote proliferation. In contrast, although TGFβ was implicated as a driver of the invasive phenotype of *TEAD1*-KO cells, *TEAD1*-KO cells failed to activate YAP1, which is in agreement with earlier studies demonstrating TEAD-dependent nuclear translocation of YAP1 ^70–72^. Importantly, despite their avid DNA-binding capacity, TEAD family members have limited transcriptional activity by themselves. Transcriptional activation depends on their ability to recruit cofactors, namely YAP1 and its paralog Taz ^59^. Although the nuclear YAP1 translocation could be mediated by the other Tef-family proteins in TEAD1-KO cells we did not find evidence supporting this in our model. Second, the transcriptomic analysis of RWPE1-α1α2-dKO and -TEAD1-KO cells revealed significant differences with α1α2-dKO cells displaying robust activation of the EMT signaling pathway while TEAD1-KO cells showed activation of myc-, TNFα and interferon pathways (data not shown). Given the similarity between the different TEAD proteins, depletion of TEAD1 might promote aberrant activation of the others potentially contributing to PCa oncogenesis. Indeed, Tef-4 (*TEAD2*) has been shown to promote TGFβ-driven EMT ^70^.

TGFβ signaling is thought to play a dualistic role in solid tumors. While it initially limits cancer growth by inhibiting cell cycle progression and inducing apoptosis at later stages of tumorigenesis TGFβ signaling promotes proliferation, invasion and drug resistance ^73^. Uncoupling of the proapoptotic signaling arm of TGFβ cascade in pretumorigenic cells is thought to convert TGFβ from a tumor suppressor into an oncogene. Moreover, TGFβ is a strong immune suppressant. Thus, autocrine TGFβ signaling has potentially strong protumorigenic effects. TGFβ is secreted in an inactive latent form that can be stored in the ECM within the tumor microenvironment. Activation of TGFβ is a complex process that can occur via multiple mechanisms many of which involve integrin functions ^74^. Interestingly, we found that cells lacking α1-integrin activated TGFβ production and secretion, but it remained mostly in inactive latent form. However, in α1α2-dKO cells lacking both collagen receptors, upregulated TGFβ was efficiently secreted and activated in autocrine manner. One potential mechanism underlying TGFβ activation could be the observed upregulation of αV-integrins in α1α2dKO cells. In particular, αVβ6- and αVβ8-integrins are critical activators of latent TGFβ ^75, 76^. Moreover, fibronectin expression is also upregulated in α1α2-dKO cells. Fibronectin is not only a ligand for αV-class integrins but it also sequestrates latent TGFβ complex and is essential for TGFβ activation ^77^. Whether an integrin switch from laminin-(α6β4) and collagen-binding (α1β1 and α2β1) integrins to RGD-binding integrins (αV-class integrins and α5β1) is indeed driving PCa tumorigenesis merits further studies. Interestingly, a bispecific antibody targeting αV- and α5β1-integrins, both of which were upregulated in α1α2-dKO cells, was shown to inhibit PCa tumorigenicity ^78^. In addition, high expression levels of αVβ5- and αVβ3 integrins have been detected in metastatic PCa ^79, 80^.

Taken together, our data identifies downregulation of α1- and α2-integrins as a risk factor in PCa. Loss or deletion of *ITGA1*/*ITGA2* locus was found in more than 2% of PCa cases but was also equally prevalent in ovarian and esophageal cancers, suggesting that the tumor suppressor functions of α1- and α2-integrins might not be limited to PCa. Loss of these collagen receptors triggered autocrine TGFβ-signaling and subsequent EMT leading to increased invasive and metastatic potential of α1/α2-integrin deficient PCa cells *in vitro* and *in vivo*. α1/α2-depleted cells also activated YAP1 signaling, that contributed to cell proliferation in these cells. Moreover, TEAD1 was identified as a key transcriptional regulator of *ITGA1* and *ITGA2* whose downregulation also contributed to PCa progression. The *TEAD1*/*ITGA1*/*ITGA2*-axis can potentially be exploited for both diagnostics purposes and for the development of more potent therapies targeting malignant forms of PCa.

## Supporting information

Cruz&Zhang_etal_Supplementary_Data_File

## ACKNOWLEDGEMENTS

Riitta Jokela is acknowledged for overall expert technical assistance, Jaana Träskelin for expert technical assistance at Biocenter Oulu Virus Core Laboratory and Dr. Veli-Pekka Ronkainen for expert assistance in microscopy at Biocenter Oulu Tissue Imaging Center. Oulu Laboratory Animal Center supported by University of Oulu is acknowledged for technical advice with mouse work. This work was funded by the Academy of Finland (251314/AM and Academy of Finland profiling program (grant #311934), Jane and Aatos Erkko foundation (AM, GHW), Sigrid Jusélius Foundation (GHW), Cancer Foundation Finland (GHW, AM) and the support funds from the Fudan University and the University of Oulu. The authors declare that they have no conflicts of interest with the contents of this article.

## AUTHOR CONTRIBUTIONS

Conceptualization: AM, GHW; Methodology: SCP, QZ, RD, TW, KZ; Validation: SCP, QZ; Formal Analysis: SCP, QZ, RD, AA, MV; Investigation: SCP, QZ, RD, JX, CP, BL, KZ; Resources: AA, MV, KZ, QZ, RD, TW; Writing – Original Draft: AM, SCP, QZ; Writing – Review & Editing: AM, GHW, SCP, QZ, RD, TW; Visualization: QZ, SCP; Supervision: AM, GHW; Funding Acquisition: AM, GHW. All authors reviewed the results and approved the final version of the manuscript.

## Abbreviations

AR: androgen receptor;
PCa: prostate cancer;
ChIP: Chromatin immunoprecipitation;
ChIP-seq: Chromatin immunoprecipitation sequencing;
RNA-seq: RNA sequencing;
GSEA: Gene set enrichment analysis;
PSA: Prostate specific antigen;
TGFβ: Transforming growth factor-β;
EMT: Epithelial-mesenchymal transition;
CEAS: cis-regulatory element annotation system.
YAP1: yes-associated protein 1;
TEAD1: Transcription enhancer factor 1

