## Supplementary material for "TEAD1 regulates ITGA1 and ITGA2 to control prostate cancer progression": Cruz&Zhang_etal_Supplementary_Data_File

#### Contents

**Supplementary Figure legends**

**Supplementary Figures 1-10**

**Supplementary Tables S1-S6**

**Supplementary Figure 1. Copy number loss/deletion of the Integrin signaling pathway components *ITGA1* and *ITGA2* associates with PCa severity.** (**a**) The meta-analysis identified 55 differentially expressed genes in the Integrin signaling pathway. *P* values determined by the Mann-Whitney U test or Pearson correlation. **(b-c)** PCa tumors with *ITGA1* (**b**) or *ITGA2* (**c**) copy number loss/del are accompanied with reduced expression levels. (**d-e**) Fraction of tumors harboring *ITGA1* (**d**) or *ITGA2* (**e**) copy number loss/del was markedly elevated in metastatic tumors compared to localized PCa. *P* values were examined by the Fisher's exact test. (**f-i**) The E2F targets and the G2M checkpoint from the Hallmark gene sets were significantly enriched in PCa tumors with *ITGA1* (**f-g**) or *ITGA2* (**h-i**) copy number loss/del vs diploid in the TCGA PCa cohort. NES, normalized enrichment score. (**j-o**) PCa patients with *ITGA1* (**j-l**) or *ITGA2* (**m-o**) copy number loss/del are associated with shorter biochemical recurrence-free survival in multiple independent PCa cohorts.

**Supplementary Figure 2. *ITGA1* and *ITGA2* are downregulated upon PCa development and progression.** (**a-h**) Box plots displaying *ITGA1* (**a-d**) or *ITGA2* (**e-h**) downregulation in human primary and metastasis PCa. *P* values determined by the Kruskal-Wallis H test or the Mann-Whitney U test. (**i-k**) Kaplan-Meier curves depicting increased risks of the biochemical recurrence (**i-j**) and overall survival (**k**) of PCa patients with lower *ITGA1* expression levels. (**l-n**) Kaplan-Meier curves depicting the associations between biochemical relapse (**l-m**) or overall survival (**n**) and *ITGA2* expression levels in the tumors of PCa patients. Patient groups were stratified by median levels *ITGA1* or *ITGA2*. The log-rank *P* values were denoted in the figures. (**o-q**) PCa patients with higher tumor stage (**o-p**) or Gleason score (**q**) were associated with downregulated *ITGA1*. (**r-s**)

*ITGA2* expression levels were decreased in PCa tumors with higher tumor stage (**r**) or Gleason score (**s**).

**Supplementary Figure 3. Immunohistochemical analysis of the PCa TMA for α1-integrin, α2-integrin and TEAD1 expression.** (**a**) Representative images of α1-integrin, α2-integrin and TEAD1 expression in normal and PCa cancer tissue. Scale bar=100µm. (**b-d**) Scoring of α1-integrin, α2-integrin and TEAD1 staining intensities in the different cell populations was done as described in Methods. The graphs show matched pairs of normal and cancer tissues from the same patients. The comparison of staining intensity was performed using two-way Anova. Asterisks indicate significance (p-value: ****<0.001).

**Supplementary Figure 4. Loss of both α1- and α2-integrins abolish cell spreading on collagen.** (**a**) RWPE1-WT, -α1KO, -α2KO and -α1α2dKO cell lysates were analyzed by western blotting for expression of α1- and α2-integrins. β-tubulin was used as a loading control. Single cell clones were picked to generate KO cell lines from parental RWPE1 cells transduced with sgRNAs targeting α1- or α2-integrins. Two independent sgRNAs were used for each target. Representative clones used in the study as biological replicates are shown. (**b**) Cells were grown in 3D Matrigel for 7 days, fixed and stained for DNA (blue) and actin (red). A representative example of a cyst with lumen, multilumen and no lumen phenotype is shown. The scale bar is 10µm. (**c**) Representative images of the indicated cell lines on collagen-coated tissue culture plates 3h, 6h and 12h after seeding. The images were captured with the IncuCyte S3 system. Scale bar=10µm. (**d**) Quantitative analysis of the percentage of spread cells in the IncuCyte S3 adhesion assay. The data shows mean ± SD. ∗ = $p < 0.05$; ∗∗ = $p < 0.01$; ∗∗∗ = $p < 0.001$.

**Supplementary Figure 5. Validation of the RWPE1 KO cell lines used for transcriptomic analysis and their correspondent top ranked pathways.** (**a-d**) Expression correlation between two biological replicates of RWPE1 control (**a**), α1α2-dKO (**b**), α1-KO (**c**) or α2-KO (**d**) RNA-seq samples, respectively. The latter three replicates expressing different sgRNAs targeting the same gene (**e-f**) Top-ranked pathways enriched in the upregulated genes form the GSEA analysis in RWPE1 cells with α1KO (**a**) or α2KO.

**Supplementary Figure 6. TGFβ pathway activation requires loss of both α1- and α2-integrins.** (**a**) Western blot analysis of latent and active TGFβ levels secreted by RWPE1-WT, -α1KO, -α2KO and -α1α2dKO cells. The data is representative of three independent experiments with similar results. (**b**) Quantification of the western blot analysis of latent and active TGFβ levels secreted by RWPE1-WT, -α1KO, -α2KO and -α1α2dKO cells. The data shows mean ±SD from three independent experiments. ∗ = $p < 0.05$; ∗∗ = $p < 0.01$; ∗∗∗ = $p < 0.001$. (**c**) RWPE1-WT, -α1KO, -α2KO and -α1α2dKO cells were grown on glass coverslips for 3 days, fixed and stained for nuclei (blue), actin (red) and YAP1 (green). Scale bar=10µm. (**d**) Quantitation of the YAP subcellular localization in the indicated RWPE1 variants. The data is representative of three independent experiments with similar results.

**Supplementary Figure 7. YAP1 is a potential transcriptional regulator of *ITGA1* and *ITGA2*.** (**a**) Quantitative PCR analysis for *YAP* levels in RWPE1-WT and RWPE1-YAP-KD cells. (**b**)YAP protein levels were assessed by Western blotting in RWPE1 with or without YAP1-knockdown (YAP KD). (**c**) Protein quantification for YAP1 levels in RWPE1 and RWPE1-YAP KD cells. (**d**) Quantitative PCR analysis for *ITGA1* and *ITGA2* levels in RWPE1-WT and RWPE1-YAP-KD cells. The data in b and c show mean ±SD from a representative experiment performed with duplicates from three independent experiments. (**e**) Quantitative PCR analysis for *ITGA1* and *ITGA2* levels in PC3-WT and PC3-YAP-KD cells. The data in b and c show mean ±SD from a representative experiment performed with duplicates from three independent experiments. (**f**) Quantitative PCR analysis for *YAP* levels in RWPE1-α1α2dKO and RWPE1-α1α2dKO+YAP-KD cells. The data in a-f show mean ±SD from a representative experiment performed with duplicates from three independent experiments. (**g**) YAP1 protein levels were assessed by Western blotting in RWPE1-α1α2dKO and RWPE1-α1α2dKO+YAP-KD cells. (**h**) Quantification of the YAP1 levels from western blots of the previous figure with three independent experiments. (**i**) Western blotting data of secreted TGFβ in RWPE1-α1α2dKO and RWPE1-α1α2dKO+YAP-KD cells. The data is representative of three independent experiments. (**j**) Proliferation of RWPE1-WT, RWPE1-α1α2dKO and RWPE1-α1α2dKO+YAP-KD cells was assessed using an XTT assay. The data shows mean ± SD from three independent experiments each performed in triplicates. $* = p < 0.05$; $** = p < 0.01$; $*** = p < 0.001$. **(k)** The Z-score sum of expression levels of *ITGA1* and *ITGA2* demonstrates positive linear correlation with the *YAP-1* expression levels in multiple human PCa tumors. *P* values examined by the Pearson's product-moment correlation test. **(l)** Genome browser representation of ChIP-seq enriched profiles of TEAD1 and YAP1 at promoters and surrounding regulatory regions of *ITGA1* and *ITGA2* in cancer cells of prostate, breast, and lung.

**Supplementary Figure 8. TEAD1 is the top-ranking gene showing the positive co-expression correlation with ITGA1 and ITGA2 in PCa tumors.** (**a-b**) *TEAD1* displayed high expression correlation with *ITGA1* (**a**) and *ITGA2* (**b**) in the MSKCC cohort of PCa tumors compared to *TEAD2*, *TEAD3* or *TEAD4*. *P* values were assessed by the two-sided Pearson's product-moment correlation test. (**c-e**) Significant positive expression correlation was identified between *ITGA1* and *ITGA2* (**c**), *ITGA1* and *TEAD1* (**d**), and *ITGA1* and *TEAD1* (**e**) in the MSKCC PCa cohort. The color bars on the right side of each figure indicate the expression of *TEAD1*, *ITGA2* and *ITGA1*, respectively. *P* values were evaluated by the Pearson's product-moment correlation test. (**f**) The top enriched motifs in the TEAD1 ChIP-seq peaks determined by HOMER software. (**g**) Genomic annotation of the input (left graph) and TEAD1 (right graph) ChIP-seq peaks in PC3 cells. Genomic features of binding peaks were visualized in pie charts, that demonstrate the different genomic features, including promoter regions within 1 kb, 1–2 kb and 2–3 kb; gene body (5′UTR, 3′UTR, exons, and introns); downstream elements and distal intergenic regions. UTR, untranslated region.

**Supplementary Figure 9. Low expression levels of *TEAD1* are associated with PCa severity.** (**a-b**) PCa tumors with *TEAD1* loss/del demonstrated significant low expression level. (**c-e**) *TEAD1* expression levels were downregulated in PCa primary (**c**, **d**) and metastatic (**e**) tumors compared to the normal prostate glands. (**f**) *TEAD1* downregulation correlated with poor prognosis of the PCa patients with biochemical recurrence.

**Supplementary Figure 10. Low levels of *ITGA1, ITGA2* and *TEAD1* correlate with tumor progression and poor prognosis in PCa patients with intermediate Gleason scores.** (**a-l**) Expression levels of *ITGA1, ITGA2, or TEAD1* cannot stratify PCa patients with lower (Gleason score ≤ 6, **a-f**) or higher risks (Gleason score ≥ 8, **g-l**). (**m**) Triple low expression level of *ITGA1*, *ITGA2* and *TEAD1* correlated with higher biochemical recurrence risk for the PCa patients. Patients were stratified by the median expression values of *ITGA1*, *ITGA2* or *TEAD1*. *P* values were examined by the log-rank test. (**n**) PCa patients with triple low expression of *ITGA1*, *ITGA2* and *TEAD1* correlated with higher PSA levels. *P* value was calculated by the two-sided Fisher's exact test.

### Cruz & Zhang et al Supplementary Figure 1

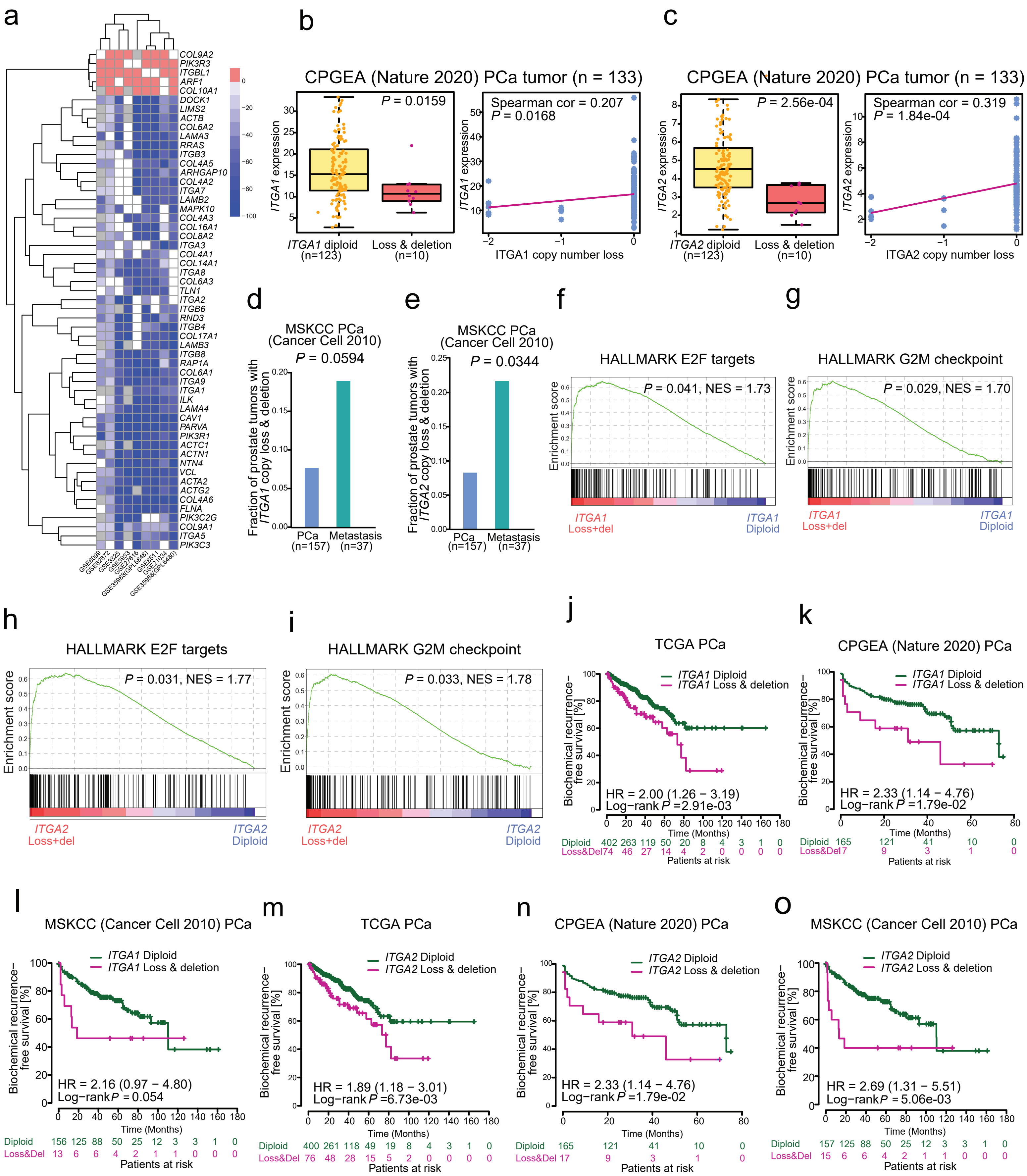

### Cruz & Zhang et al  Supplementary Figure 2

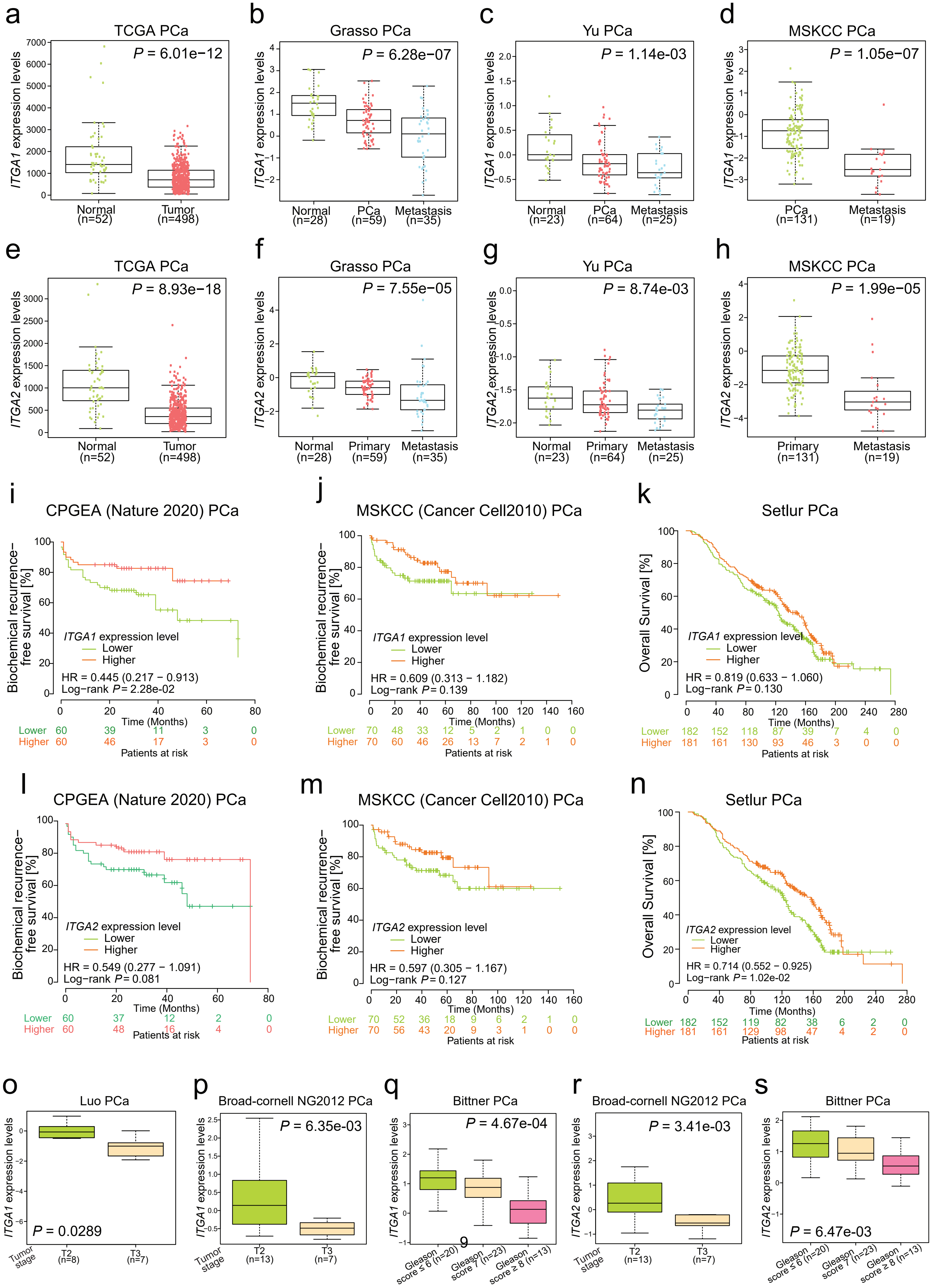

**Cruz & Zhang et al  Supplementary Figure 3**

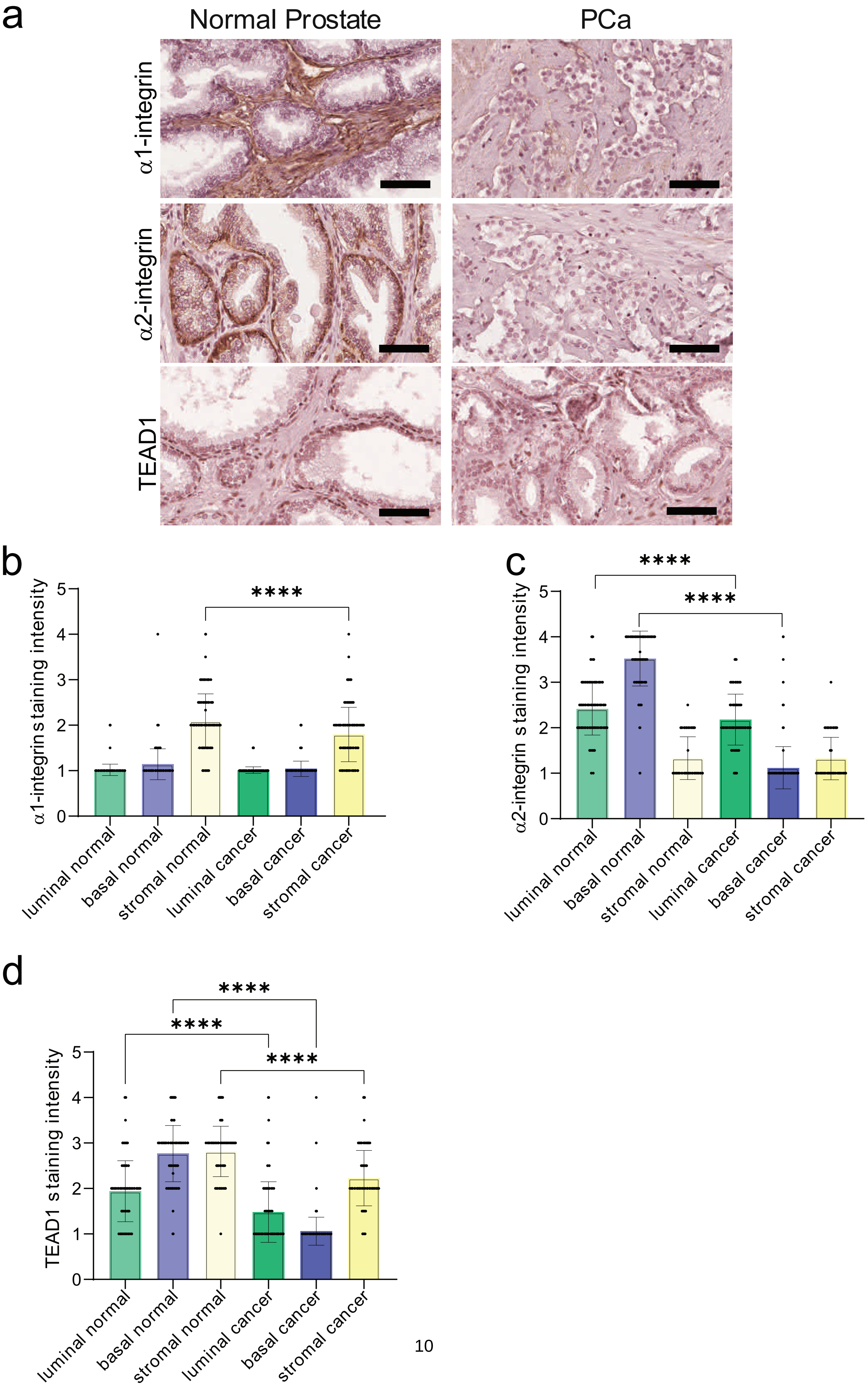

### Cruz & Zhang et al  Supplementary Figure 4

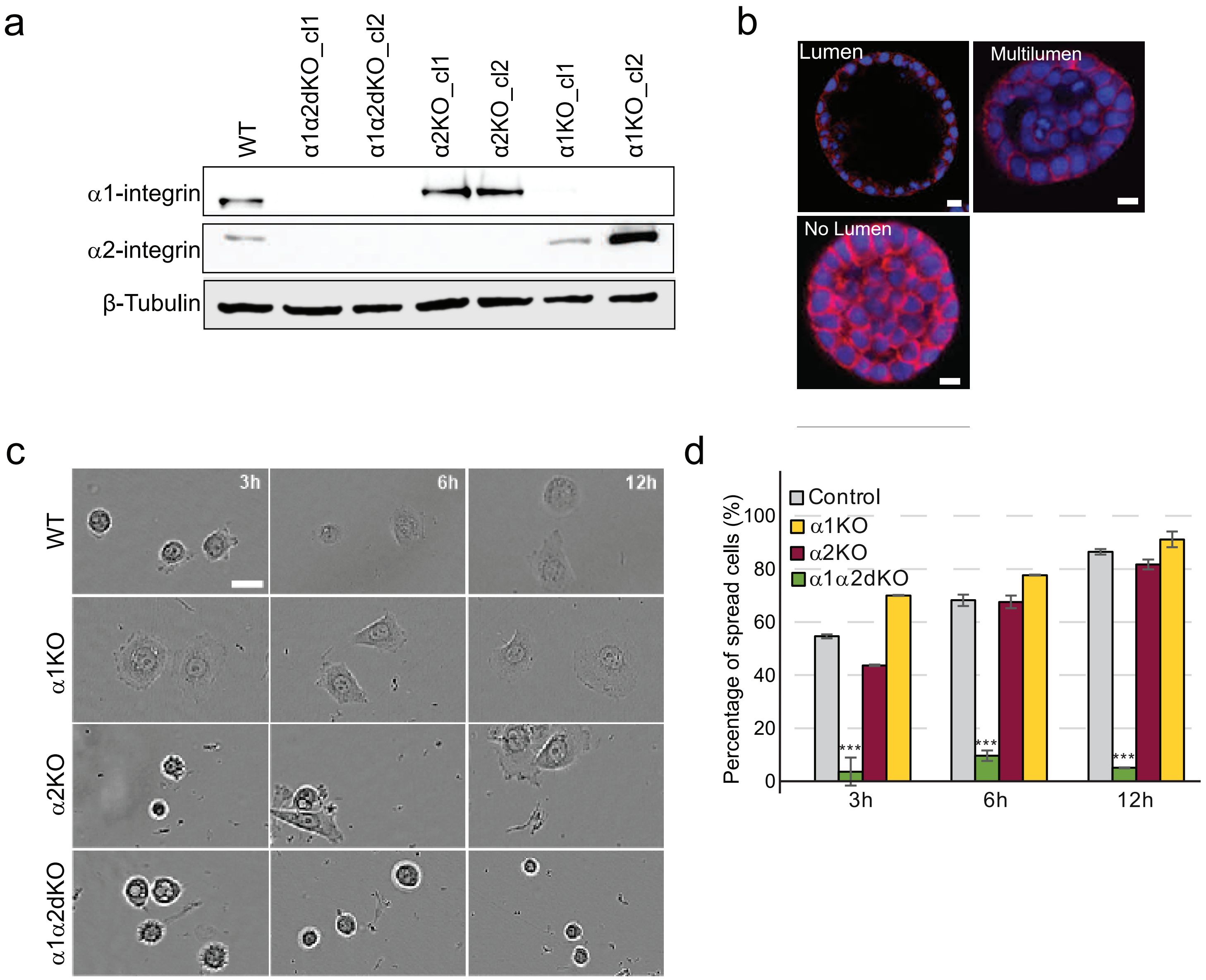

**Cruz & Zhang et al  Supplementary Figure 5**

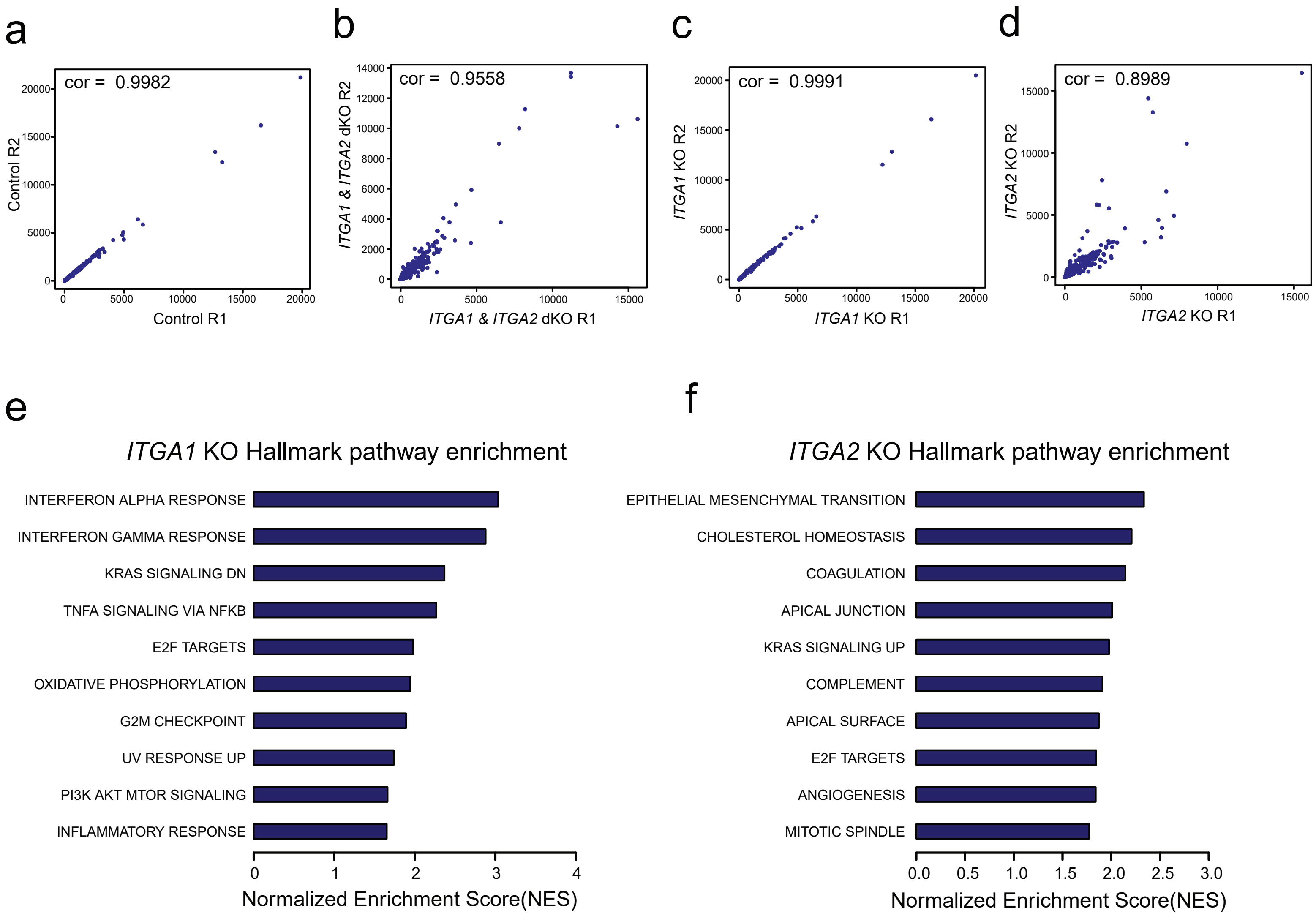

Cruz & Zhang et al Supplementary Figure 6

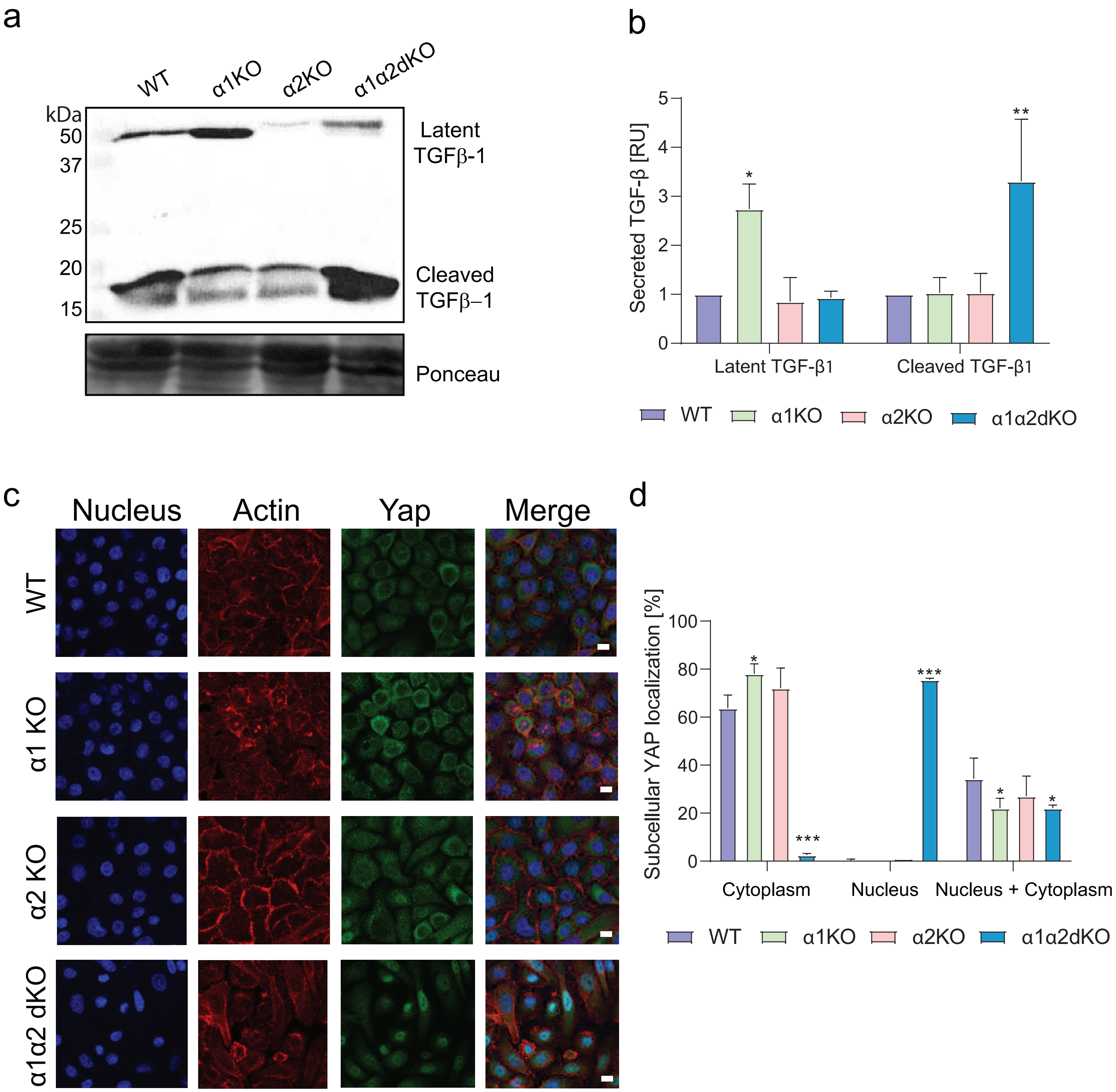

### Cruz & Zhang et al Supplementary Figure 7

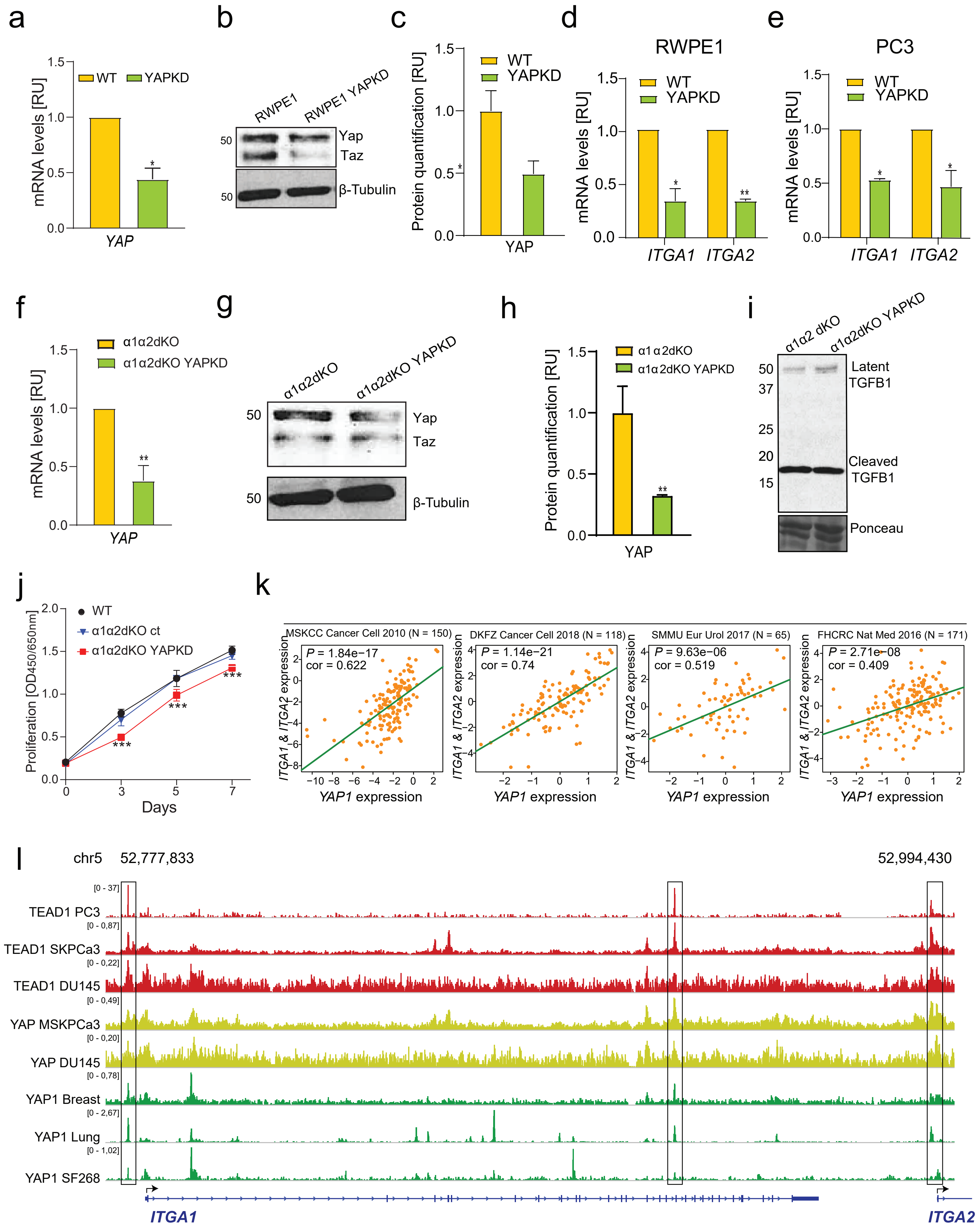

### Cruz & Zhang et al Supplementary Figure 8

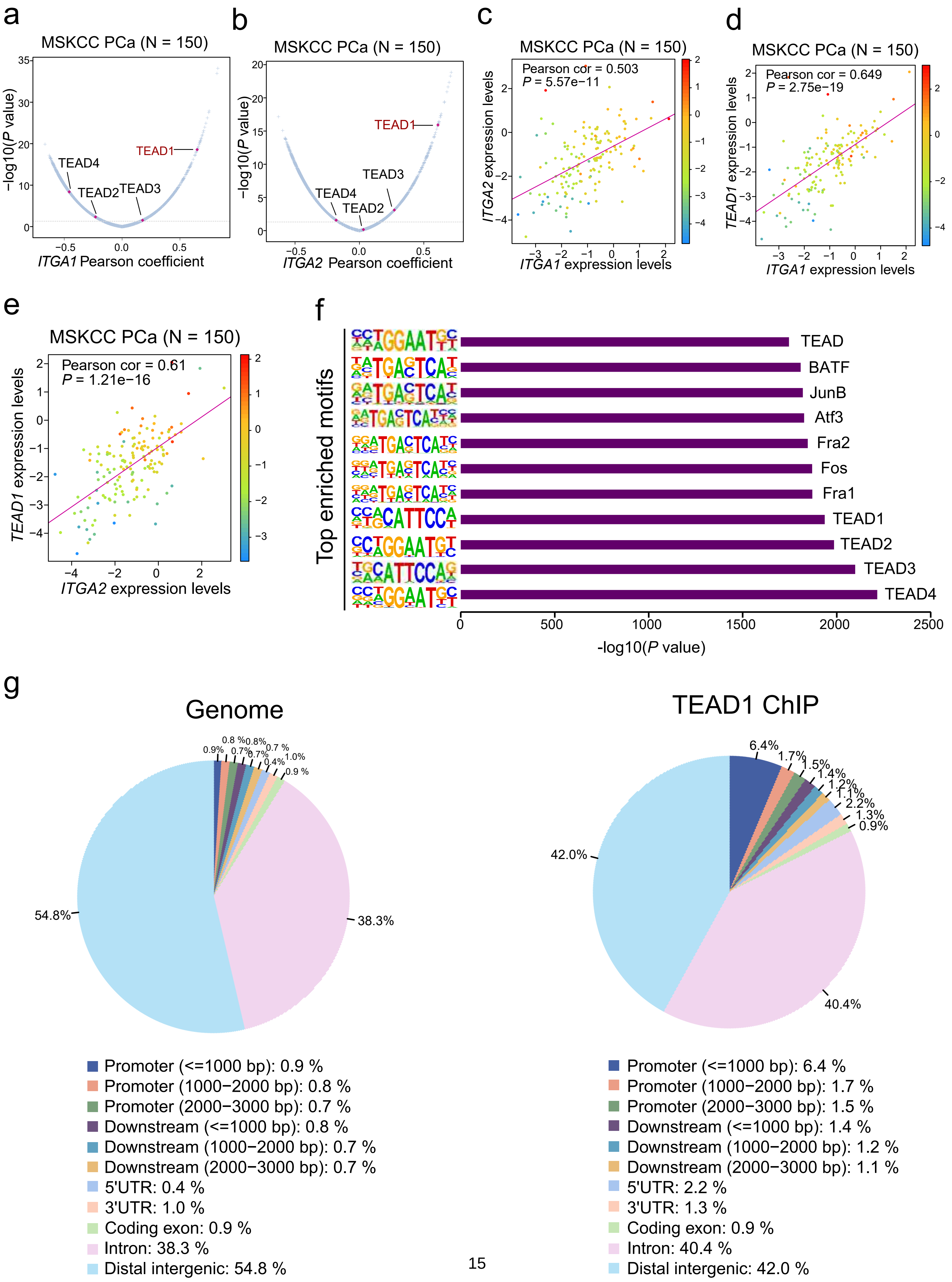

**Cruz & Zhang et al Supplementary Figure 9**

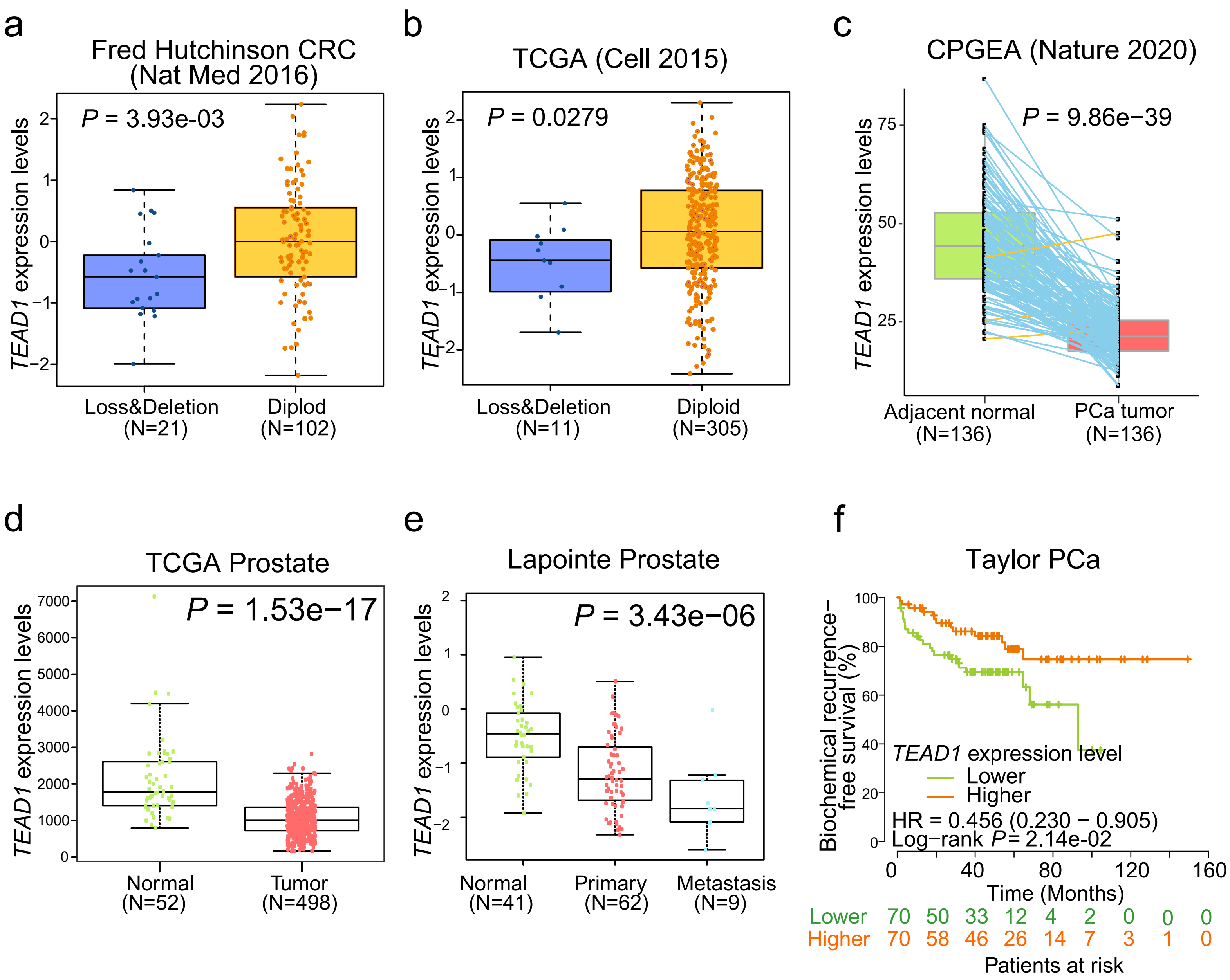

### Cruz & Zhang et al Supplementary Figure 10

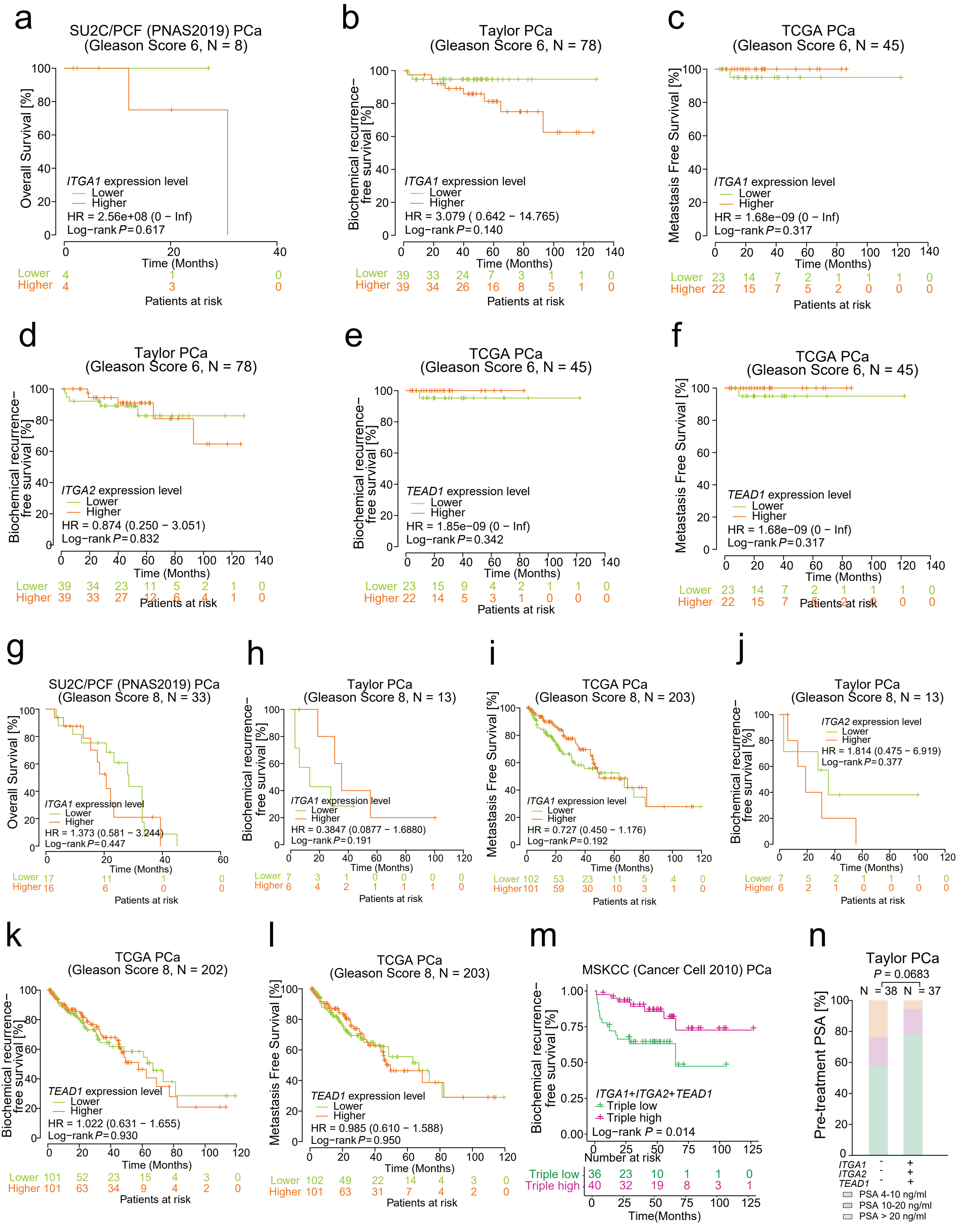

#### Supplementary Tables S1-S6

##### Table S1 List of antibodies used in this study

| | Name | Company | Cat. No | Dilution for WB | Dilution for ICC/IHC |
|---|---|---|---|---|---|
| 1 | anti α1-integrin | R&D Systems | AF5676 | 1:200 | |
| 2 | anti α1-integrin | Abcam | Ab243032 | | 1:1000 |
| 3 | anti α2-integrin | Santa Cruz | sc-9089 | 1:500 | |
| 4 | anti α2-integrin | Abcam | Ab181548 | | 1:50 |
| 5 | anti-TEAD1 | BD Transduction Laboratories | 610922 | 1:500 | |
| 6 | anti-TEAD1 | Cell Signaling Technology | 12292S | | 1:75 |
| 7 | anti-TGFβ1 | Abcam | Ab179695 | 1:1000 | |
| 8 | anti-Yap-1/Taz | Cell Signaling | 8418 | 1:500 | |
| 9 | anti-β-tubulin | Sigma | T4026 | 1:7500 | |

#### Table S2 Target sequences of CRISPR/Cas9

| **Name** | **Gene** | **Target region** | **Oligo sequence 5' to 3'** |
|---|---|---|---|
| *ITGA1-KO1-F* | *ITGA1* | CDS exon 1 | CACCGAATGACTTTCAGCGGCCCGG |
| *ITGA1-KO1-R* | *ITGA1* | CDS exon 1 | AAACCCGGGCCGCTGAAAGTCATTC |
| *ITGA1-KO2-F* | *ITGA1* | CDS exon 2 | CACCGCTTATTGGTTCTCCGTTAGT |
| *ITGA1-KO2-R* | *ITGA1* | CDS exon 2 | AAACACTAACGGAGAACCAATAAGC |
| *ITGA2-KO1-F* | *ITGA2* | CDS exon 2 | CACCGTTCTGGGAGACCAACATTGT |
| *ITGA2-KO1-R* | *ITGA2* | CDS exon 2 | AAACACAATGTTGGTCTCCCAGAAC |
| *ITGA2-KO2-F* | *ITGA2* | CDS exon 3 | CACCGGGTCCTTCAAGTGAACAGTT |
| *ITGA2-KO2-R* | *ITGA2* | CDS exon 3 | AAACAACTGTTCACTTGAAGGACCC |
| *TEAD1-KO1-F* | *TEAD1* | CDS exon 3 | CACCGCTATCTATCCACCATGTGGG |
| *TEAD1-KO1-R* | *TEAD1* | CDS exon 3 | AAACCCCACATGGTGGATAGATAGC |
| *TEAD1-KO2-F* | *TEAD1* | CDS exon 7 | CACCGTGGCCGGGAATGATTCAAAG |
| *TEAD1-KO2-R* | *TEAD1* | CDS exon 7 | AAACACCGGCCCTTACTAAGTTTGC |

#### Table S3 List of Target sequence of shRNA

| **Name** | **Gene** | **Target sequence 5' to 3'** |
|---|---|---|
| *shYAP1-1* | *YAP-1* | GACCAATAGCTCAGATCCTTT |
| *shYAP1-2* | *YAP-1* | CAGGTGATACTATCAACCAAA |

#### Table S4 List of qPCR primers

| **Gene** | **Sequence 5’-3’** |
|---|---|
| *ITGA1* | F: CTGGACATAGTCATAGTGCTGGA<br>R: ACCTGTGTCTGTTTAGGACCA |
| *ITGA2* | F: CACCGGGTCCTTCAAGTGAACAGTT<br>R: AAACAACTGTTCACTTGAAGGACCC |
| *ITGA5* | F: GCCTGTGGAGTACAAGTCCTT<br>R: AATTCGGGTGAAGTTATCTGTGG |
| *ITGAV* | F: AACTCAAGCAAAAGGGAGCA<br>R: TCATGTTCTTGGAGTGACTTGG |
| *SLUG* | F: GATCTGCCAGACGCGAACTC<br>R: GGCAACCAGACAACCGACAT |
| *TWIST* | F: GCCGGAGACCTAGATGTCATTG<br>R: CCCACGCCCTGTTTCTTTGA |
| *SNAII* | F: CAGTGCCTCGACCACTATGC<br>R: GCAGCTCGCTGTAGTTAGGC |
| *CDH1* | F: CACCGGTCGACAAAGGACAG<br>R: TCCAGAAACGGAGGCCTGAT |
| *CDH2* | F: TCAACCCCATACACCAGCCT<br>R: AGTCGATTGGTTTGACCACGG |
| *YAP1* | F: GAACTCGGCTTCAGGTCCTC<br>R: GGTTCATGGCAAAACGAGGG |
| *GAPDH* | F: AACAGCGACACCCATCCTC<br>R: CATACCAGGAAATGAGCTTGACAA |

#### Table S5 Primer sequences of ChIP-qPCR

| | |
|---|---|
| ITGA1pro-TEAD1-71F | CAGCTAAAATGTTACTGCTCATCA |
| ITGA1pro-TEAD1-71R | CAGGAATGTTCAATTCGTGGCA |
| ITGA1-TEAD1-73F | TCATCAGGCTTTTGCTTCCCT |
| ITGA1-TEAD1-73R | CTGAAACCTCCACATTCCCCT |
| ITGA2pro-TEAD1-97F | CCCAAACACAGGTCTGTTCC |
| ITGA2pro-TEAD1-97R | AAGGAAGTCAGCCAGGTTTCA |
| SLC1A5 Posi -TEAD1-119F | CCCTGAGGCATTGTGGGTTC |
| SLC1A5 Posi -TEAD1-119R | TGCAAGCTGTCCAGGGTATT |

#### Table S6 Clinicopathological characteristics of patients included in the TMA analysis.

| **Parameter** | **Number of cases in the group** (Total 242) |
|---|---|
| **Age at the time of surgery:** | |
| **<55** | 37 |
| **55-65** | 144 |
| **>65** | 61 |
| **Tumor stage:** | |
| **T2** | 173 |
| **T3** | 67 |
| **T4** | 2 |
